# The TOG protein Stu2 is regulated by acetylation

**DOI:** 10.1101/2021.12.31.474666

**Authors:** Matt Greenlee, Braden Witt, Jeremy Sabo, Savannah Morris, Rita K. Miller

## Abstract

Stu2 in *S. cerevisiae* is a member of the XMAP215/Dis1/Alp14/Msps/CKAP5/ch-TOG family of MAPs and has multiple functions in controlling microtubules, including microtubule polymerization, microtubule depolymerization, linking chromosomes to the kinetochore, and assembly of γ-TuSCs at the SPB. Whereas phosphorylation has been shown to be critical for Stu2 localization at the kinetochore, other regulatory mechanisms that control Stu2 function are still poorly understood. Here, we show that a novel form of Stu2 regulation occurs through the acetylation of three lysine residues at K252, K469, and K870, which are located in three distinct domains of Stu2. Alteration of acetylation through acetyl-mimetic and acetyl-blocking mutations did not impact the essential function of Stu2. Instead, these mutations lead to both positive and negative changes in chromosome stability, as well as changes in resistance to the microtubule depolymerization drug, benomyl. In agreement with our *in silico* modeling, several acetylation-mimetic mutants displayed increased interactions with γ-tubulin. Taken together, these data suggest that Stu2 acetylation can govern multiple Stu2 functions in both a positive and negative manner, including chromosome stability and interactions at the SPB.

## INTRODUCTION

Stu2 is a member of the XMAP215/Dis1/CKAP5/ch-TOG family of MAPs that has multiple microtubule-dependent functions. Stu2 is well known for its role as a microtubule polymerase (Al-Bassam et al., 2006) (Ayaz et al., 2014) (Brouhard et al., 2008) (Podolski et al., 2014) (Wang & Huffaker, 1997) reviewed in (Al-Bassam & Chang, 2011) (Al-Bassam & Chang, 2011). Stu2 promotes microtubule elongation through interactions with αβ-tubulin heterodimers via two TOG domains (Al-Bassam et al., 2007) (Al-Bassam et al., 2006) (Ayaz et al., 2014) (Ayaz et al., 2012) (Slep & Vale, 2007) and interacts with the microtubule lattice through a basic microtubule binding (MT binding) domain (Al-Bassam et al., 2006) (Wang & Huffaker, 1997). To carry out microtubule polymerization, Stu2 is also thought to undergo significant conformational changes throughout the tubulin binding and microtubule polymerization processes (Al-Bassam & Chang, 2011) (Al-Bassam et al., 2006) (Ayaz et al., 2014) (Nithianantham et al., 2018). Stu2 also interacts with other microtubule-associated proteins, such as Bik1, Bim1, Ndc80, and Spc72 through its c-terminal MAP interacting domain (Usui et al., 2003) (Aravamudhan et al., 2014) (Wolyniak et al., 2006) (Miller et al., 2016) (Miller et al., 2019).

While Stu2 is perhaps best known for contributing to microtubule stability and growth, it has also been shown to initiate microtubule depolymerization (Al-Bassam & Chang, 2011) (Al-Bassam et al., 2006) (Brouhard et al., 2008) (Humphrey et al., 2018) (Shirasu-Hiza et al., 2003) (van Breugel et al., 2003). Stu2 also functions at the spindle pole body (SPB), where interactions with Spc72 facilitate microtubule nucleation by promoting oligomerization of γ-TuSC assemblies (Chen et al., 1998) (Gunzelmann et al., 2018) (Thawani et al., 2018) (Usui et al., 2003). Additionally, interactions between Stu2 TOG, MT binding, and c-terminal MAP interacting domains and Tub4 (γ-tubulin), Spc97, and Spc72 respectfully all correlate with MT nucleation in the presence of Spc72 and tubulin with or without purified γ-TuSC (Gunzelmann et al., 2018). The γ-TuSC complex is thought to have an open form that is inactive and a closed form that actively nucleates microtubules. These two γ-TuSC states provide a structural basis for its role in MT nucleation (Greenberg et al., 2016).

Stu2 has multiple functions at the kinetochore. It is important for attachment of kinetochore microtubules (K-fibers) to the outer plaque of the kinetochore, where it also aids in the selection of correctly oriented kinetochore attachments (Aravamudhan et al., 2014). Stu2 accomplishes this task by serving as a mechano-sensor while bound to the Ndc80 complex through its c-terminal MAP binding domain (Miller et al., 2016) (Zahm et al., 2021). Levels of tension associated with correct bi-polar microtubule attachments to sister chromatids initiate Stu2 mediated microtubule plus-end depolymerization and subsequent Dam1 complex capture to secure optimal kinetochore K-fiber interactions (Asbury et al., 2006) (Humphrey et al., 2018) (Miller et al., 2016).

However, many of the regulatory mechanisms that control Stu2 function during mitosis and meiosis are poorly understood. Most recently, a Cdk1 phosphorylation site found in the basic MT binding domain of Stu2 was shown to promote Stu2 localization to the kinetochore (Humphrey et al., 2018). S603 phosphorylation of Stu2 neutralizes the basic MT binding patch and enables Stu2 to more readily concentrate at kinetochores through interactions with other MAPs (Humphrey et al., 2018). While phosphorylation reduces interactions with the MT lattice, the TOG1, the coiled coil Stu2 dimerization, and the C-terminal MAP interacting domains are also important for Stu2 localization to the kinetochore (Miller et al., 2019). Phosphorylation of other XMAP215 family members has also been reported to control their association with the microtubule lattice; in *S. pombe* CDK mediated phosphorylation was found in Alp14 (Aoki et al., 2006) and Dis1 proteins (Okada et al., 2014), and Msps phosphorylation was found in *D. melanogaster* (Trogden & Rogers, 2015). While phosphorylation of its MT-binding domain regulates Stu2 association with the microtubule lattice (Humphrey et al., 2018) (Okada et al., 2014) (Trogden & Rogers, 2015), the role that other posttranslational modifications such as sumoylation play in governing Stu2 localization and functions remain elusive (Greenlee et al., 2018).

In this report, we investigate the relationship of Stu2 and acetylation, a post-translational modification that nullifies the positive charges present on lysine side chains. We identify acetylation as a novel of Stu2 modification and show that acetylation of Stu2 occurs on three lysine residues, K252, K469, and K870. These residues are located in the TOG1, TOG2, and the MAP interacting domains, respectively. Mutational analysis revealed distinct functions associated with the three different sites. An analysis of single, double, and triple mutants suggests a hierarchy of intragenetic epistatic regulation for Stu2 acetylation at the SPB and kinetochore. These findings suggest a novel mechanism by which acetylation regulates multiple functions of Stu2.

## RESULTS

### Stu2 is acetylated at lysines 252, 469, and 870

As Stu2 has been shown to be modified by phosphorylation and sumoylation, we asked whether Stu2 could be regulated by other types of modifications, such as acetylation. For this, we immunoprecipitated HA-tagged or untagged Stu2 from logarithmically growing cultures treated with the deacetylation inhibitor Trichostatin A (TSA). The precipitate was then immuno-blotted with anti-HA to detect Stu2 and anti-AcK to detect acetylated lysines. As shown in Figure 1A, bands with a molecular weight of approximately 115 kDa were observed in both the anti-HA and anti-AcK immunoblots. Importantly, this reactivity was absent in extracts containing untagged Stu2. This finding suggests that Stu2 is acetylated.

**Figure 1.**
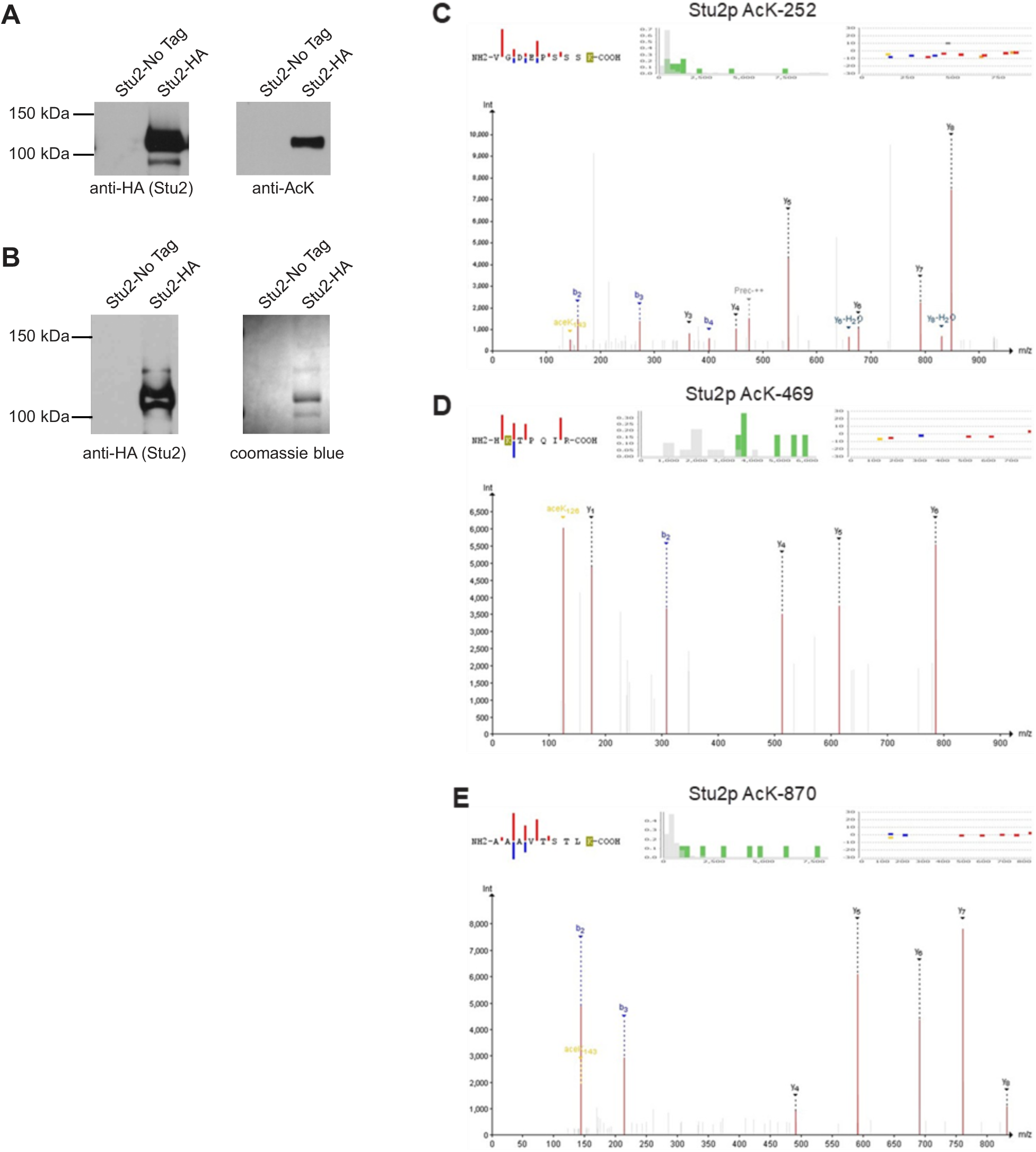
Stu2 is acetylated at lysine 252, 469, and 870. (A) Yeast extracts containing Stu2-HA (yRM12358) or untagged Stu2 (yRM12359) were immunoprecipitated with anti-HA magnetic beads and immunoblotted with mouse anti-HA and rabbit anti- AcK. (B) To identify protein bands for analysis by ms/ms mass spectrometry, Stu2-HA was immunoprecipitated with anti-HA magnetic beads. Pull-downs were separated by SDS-PAGE and either immunoblotted with mouse-anti HA to identify the Stu2-reactive bands or stained with coomassie blue R-250, which were excised. Tandem mass spectrometry identified three putatively acetylated lysines residues; (C) 252, (D) 469, and (E) 870. Mass spectra were analyzed by the six search engines; Comet, MS Amanda, MS- GF+, Myri-Match, OMSSA, and X!Tandem, using SearchGUI. SearchGUI outputs were then combined using PeptideShaker for interpretation, statistical analysis, and viewing. For the spectra shown, mass deviations of 2.19 ppm (C), 0.53 ppm (D) and 0.47 ppm (E) were observed. For each site, all six search engines identified these three lysines as the most probable acetylated peptide.

To identify specific acetylated residues, we employed a mass-spectrometry strategy. Stu2-HA was immunoprecipitated from whole cell extracts prepared from logarithmically growing cultures. The bands corresponding to Stu2 were excised from coomassie blue stained gels and prepared for tandem mass spectrometry (MS/MS) (see Materials and Methods) (Figure 1B). This analysis identified 7650 spectra corresponding to Stu2. Of these, 59 showed a 42 Da increase in mass that is characteristic of acetylation, with an error of less than 10ppm. Thirty- two of the spectra corresponded to peptides with an acetyl-modification of lysine 252. Nine spectra corresponded to acetylation of lysine 469, and 18 corresponded to acetylation of lysine 870. Nine spectra of AcK 252 and three spectra of AcK 870 had a confidence level of 100%.

Eight of the nine spectra obtained for AcK469 had a confidence score of 90% or higher. Many of these putative acetylated peptides also contained immonium reporter ions that are diagnostic for acetylated lysine (Figure 1C-E). The confidence scores for these and additional lower- confidence spectra peptides are shown in Table S1. Notably, the acetylation sites in this ms/ms analysis were identified in the absence of inhibitors or strains with compromised KDACs, limiting the possibility that they were experimentally induced. Combined, these data suggest that Stu2 is a *bona fide* multi-acetylated protein.

**Table 1.**
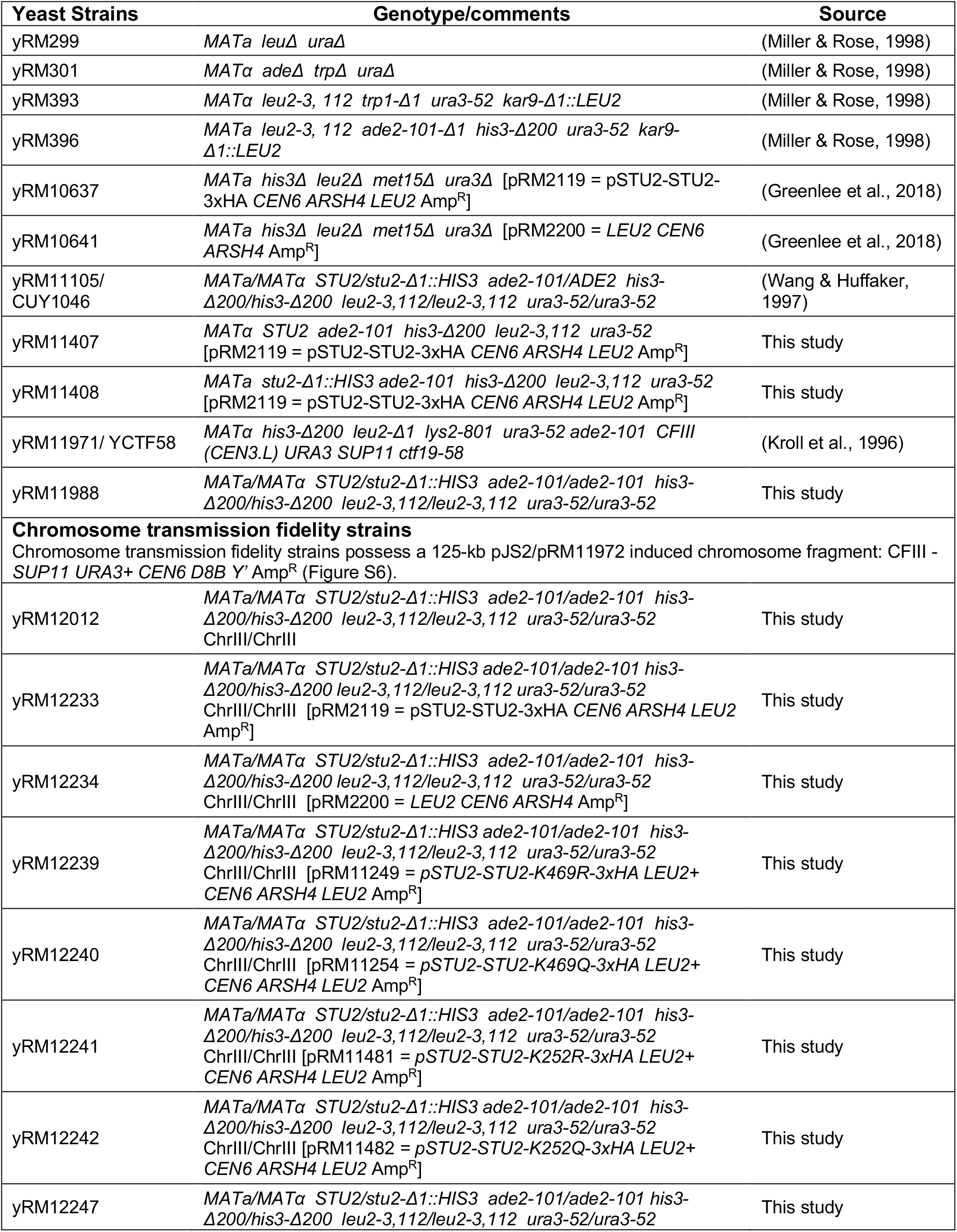
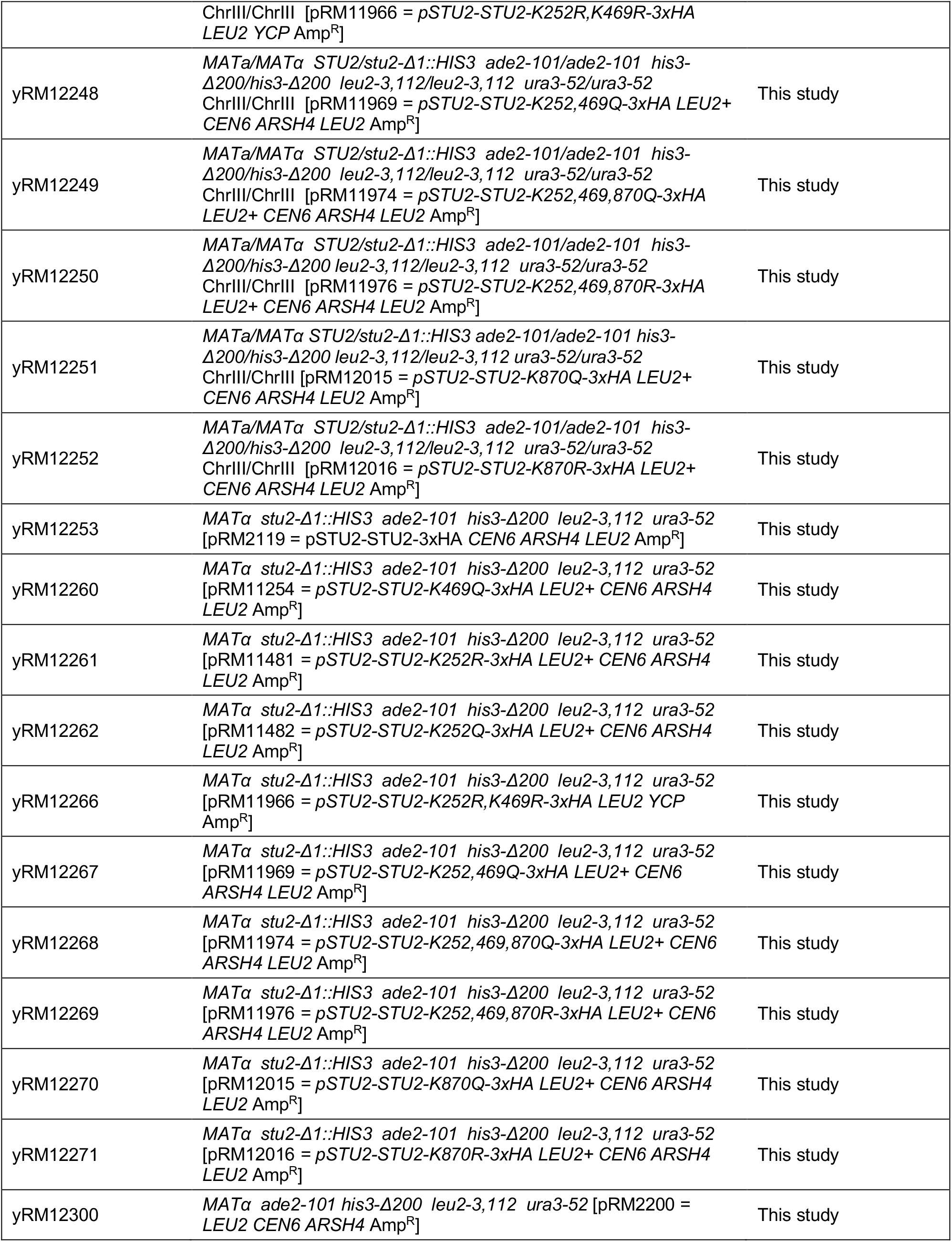
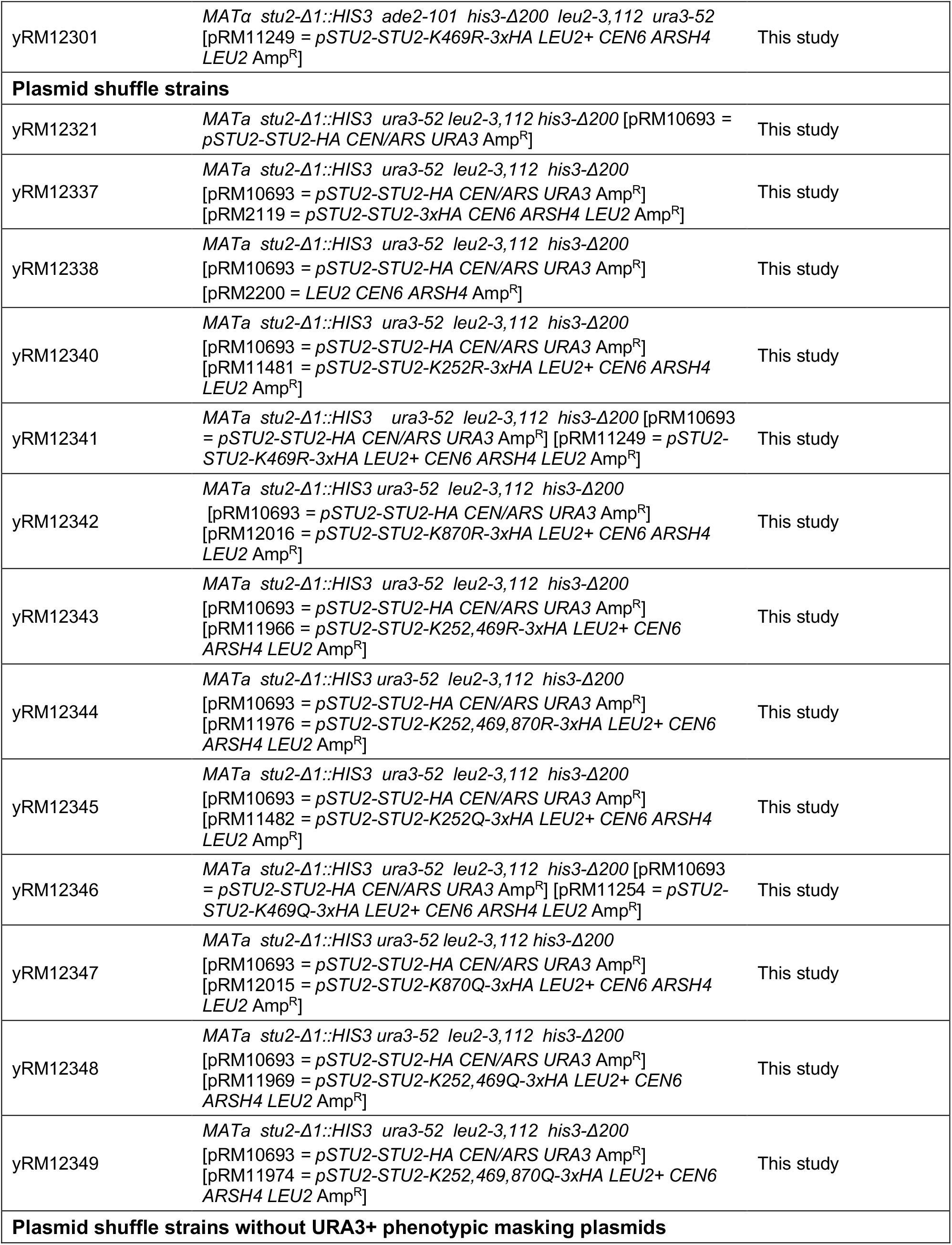
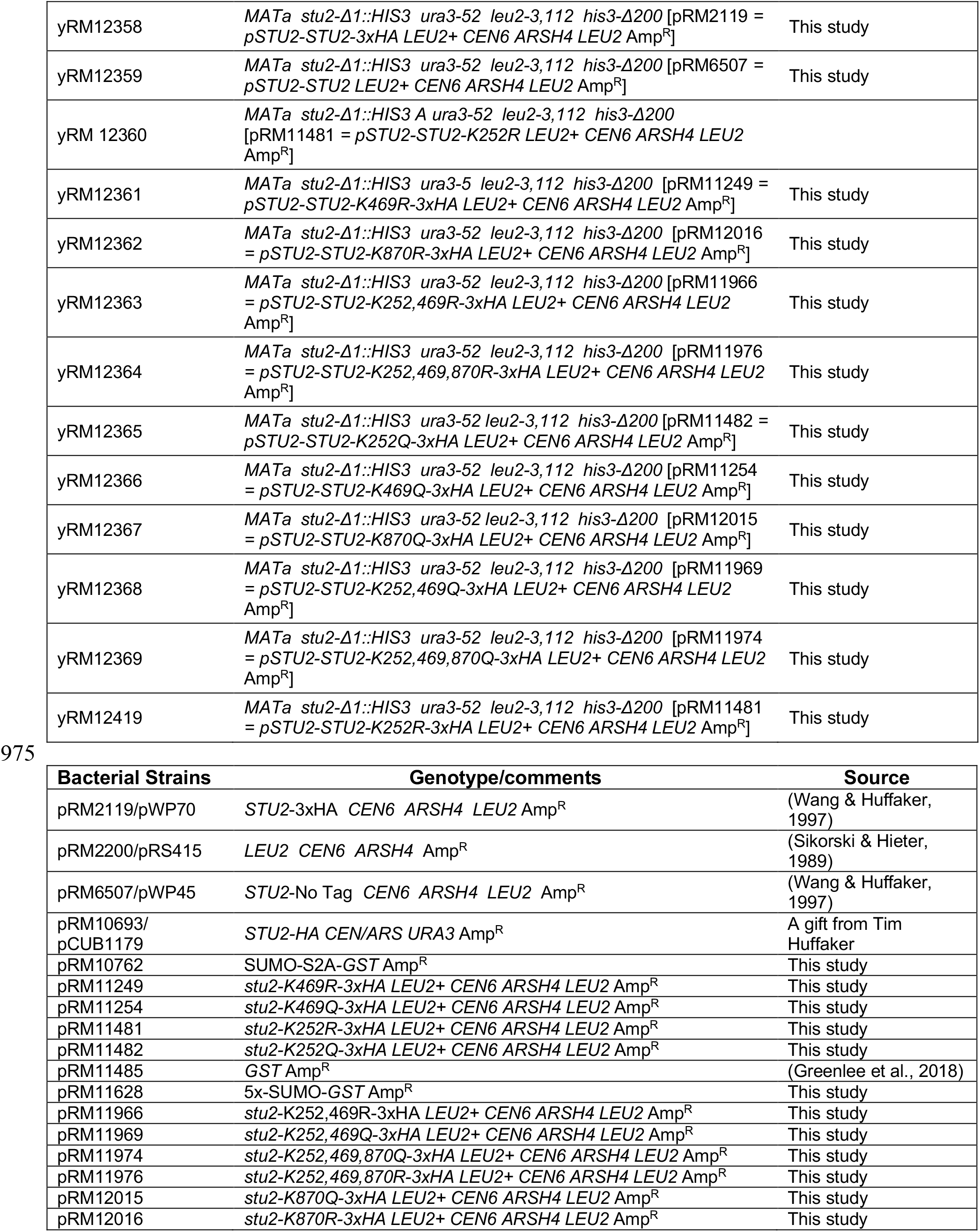
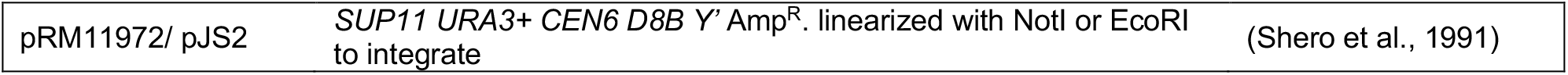
Yeast and bacteria strains.

### Acetylated lysines are found in multiple domains of Stu2

To gain insight into how lysine acetylation influences Stu2 function, we mapped each acetylated lysine to a composite of predicted and actual crystal structures of Stu2 using the structural model program I-TASSER. The I-TASSER depiction of the Stu2 TOG domains (Figure 2B) closely resembled published crystal structures of TOG domains bound to αβ-tubulin heterodimers, as the coordinates from the pdb files 4FFB and 4U3J were used (Ayaz et al., 2012) (Ayaz et al., 2014). From this model, all the acetylated lysines are predicted to be solvent exposed. The acetyl-lysine K252 lies between α-helices 14 and 15, a region that comprises a flexible linker connecting TOG1 and TOG2 domains. Lysine 469 lies between α-helices 10 and 11 of the TOG2 domain where it interacts with β-tubulin within the TOG2-tubulin binding pocket. K870 lies in the MAP interacting domain, a domain that interacts with Bik1, Bim1, Ndc80, and Spc72 (Usui et al., 2003) (Wolyniak et al., 2006) (Aravamudhan et al., 2014) (Zahm et al., 2021).

**Figure 2.**
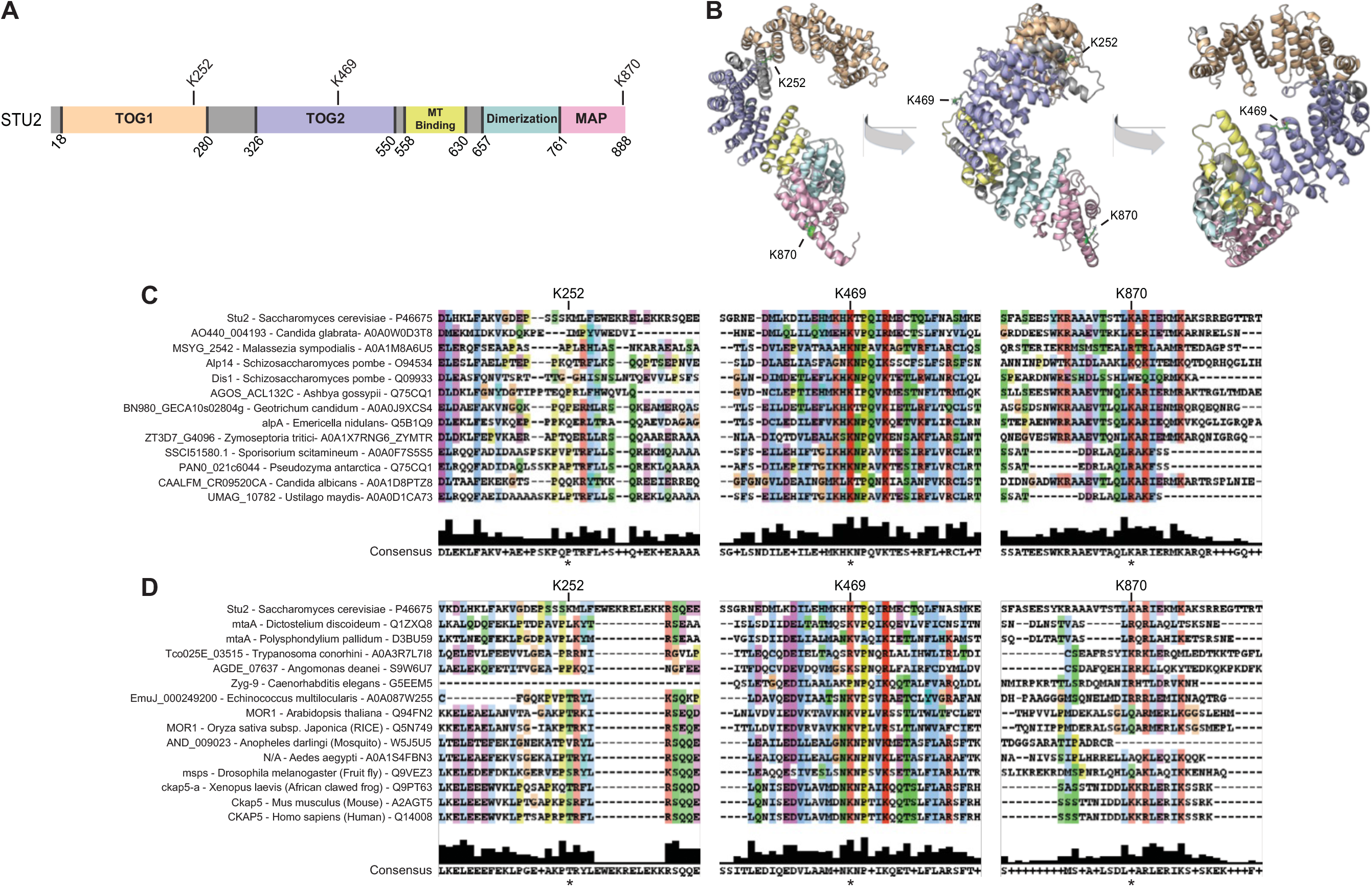
Stu2 lysines 469 and 870 are evolutionarily conserved. (A) The location of acetyl-lysines are indicated relative to distinct Stu2 domains and (B) are illustrated in a Stu2 structure predicted by I-TASSER. Clustal protein alignments are shown for members of the Stu2/XMAP215 family of microtubule polymerases for members across (C) eukaryota and (D) fungi. acetylation site may reflect diversification of the XMAP215 family and their interaction with different classes of microtubule associated proteins. Notably, two insects (*A. Aegypti, and D. melanogaster*) and two trypanosomatids (*T. conorhini, and A. deanei*) possessed a lysine at position 872, which might represent an alternative acetylation site.

To assess if the three acetyl-lysines were evolutionarily conserved amongst the Stu2/XMAP215 family of proteins, we aligned XMAP215 sequences of budding, fission, anamorphic, and filamentous fungi (Figure 2C), as well as eukaryotic organisms of varying complexity (Figure 2D). Of the three lysines, lysine 252 was the least conserved and lies in an intrinsically unstructured loop between TOG domains (Ayaz et al., 2012). It was found only in the budding yeast *S. cerevisiae* among the twenty-seven organisms analyzed. In contrast, lysine 469 was highly conserved across eukaryotes, including all fungal species analyzed. Lysine 870 was conserved in eight of the thirteen fungal and four additional eukaryotic XMAP215 proteins, including trypanosomes, frogs, mice, and humans (also see Figures 1 and 4 in Zahm *et al*., 2021 for additional alignments of this region). The lower rate of conservation for Stu2’s C-terminal

### Mutations in Stu2 acetylation sites do not disrupt the essential function of Stu2

To investigate the function of these acetylation sites, we mutated lysines 252, 469, and 870 to arginine to prevent acetylation while still maintaining their positive charge. The lysines were also substituted with glutamine to mimic acetylation. Several combinations of double and triple mutations were also constructed. Lysines 252 and 469 both reside in or near the two TOG domains. To eliminate acetylation within the TOG domain, these lysines were mutated as a pair to either arginine or glutamine. This generated the double mutants, 2KR and 2KQ. To generate the triple mutants, lysines 252, 469, and 870 were all mutated to either arginine (3KR) or glutamine (3KQ). As observed from the analysis of Stu2 in WCE prepared from logarithmically growing cultures, the 3KR and the 3KQ routinely displayed steady-state abundance levels equivalent to that of WT of Stu2 when normalized to the Pgk1 loading control (Figure 3A and Figure 3B, Figure S1A-C). Similarly, the single K-to-R and K-to-Q mutants, also displayed nearly equivalent expression levels (Figure 4A,B). This suggests that in logarithmically-growing cultures these acetylation site mutants have little or no substantial effect on Stu2 abundance.

**Figure 3.**
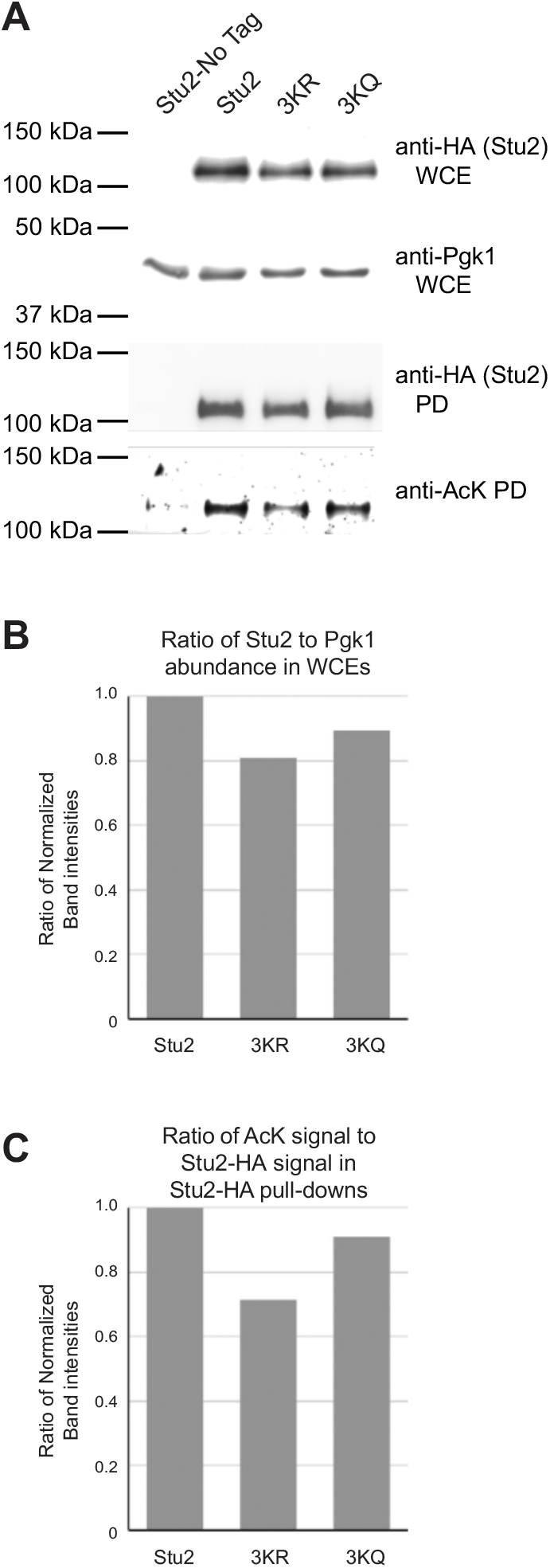
The Stu2-3K mutants display a moderate reduction in acetylation. Stu2-3K mutants with mutation of lysines 252, 469, and 870 in HA epitope- tagged strains were analyzed for acetylation by western blot. (A) Cells expressing Stu2-HA (yRM12358), untagged Stu2 (yRM12359), Stu2- 3KR-HA (yRM12364), or Stu2-3KQ-HA (yRM12369) were immunoprecipitated and western blotted with anti-HA and anti-AcK to evaluate Stu2 acetylation. Band densitometry was performed to quantify reductions in Stu2 acetylation. (B) The ratio of Stu2 present in WCEs relative to the Pgk1 loading control was evaluated to determine differences in Stu2 steady state abundance in logarithmically growing cell cultures using NIH ImageJ. (C) The ratio of anti-AcK to Stu2-HA signal from immunoprecipitations was determined to evaluate reductions in acetylation for the Stu2-3K mutants compared to wild type. In bioreplicates of this experiment, the amount of TSA used also affected the intensity of anti-acetyl-lysine reactivity with the Stu2 band (data not shown).

**Figure 4.**
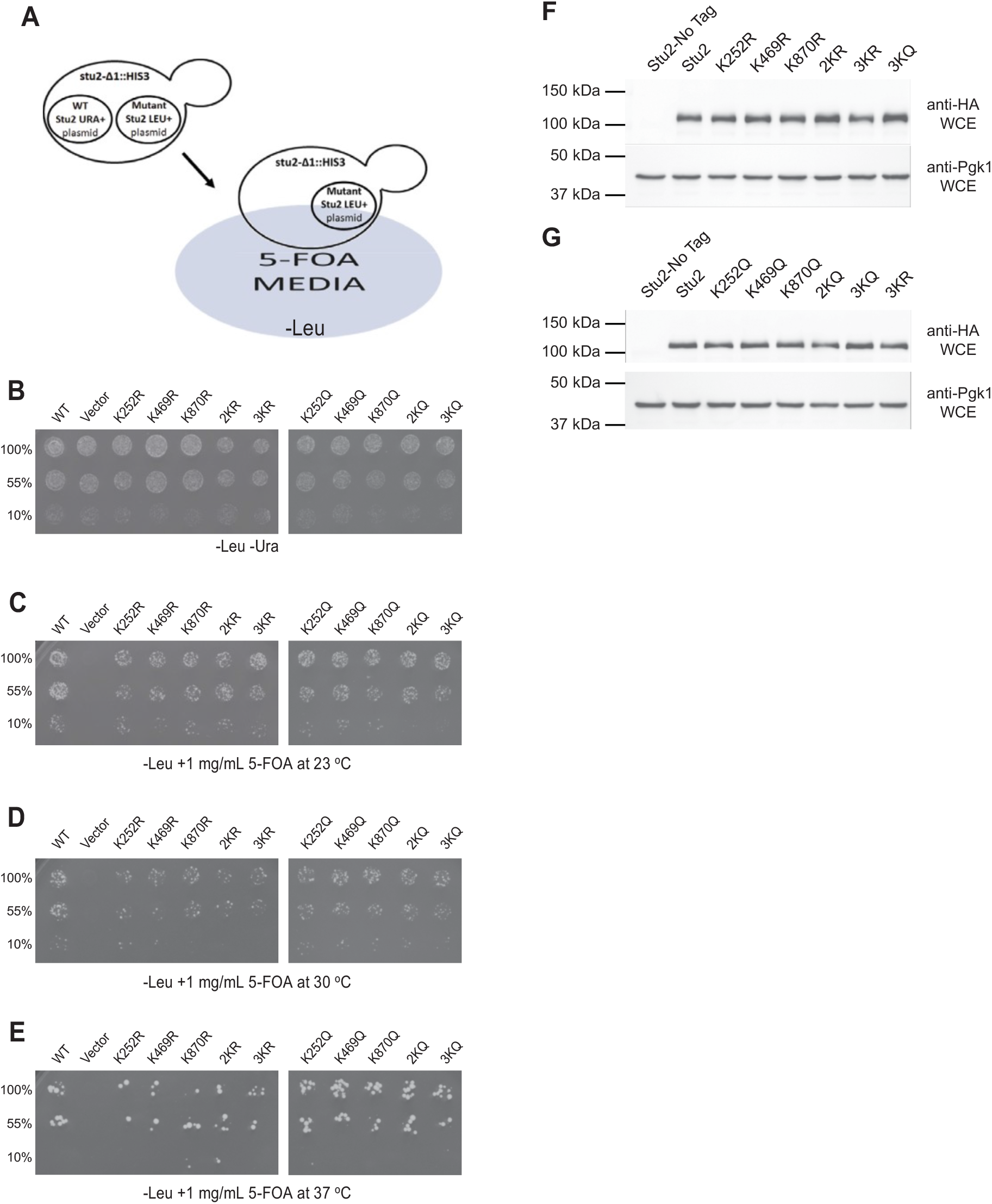
Plasmid shuffle complementation assays demonstrate Stu2 acetyl state mimetic mutants are functional. (A) A plasmid shuffle assay was used to detect phenotypes from acetyl-lysine mimicking (K-to-R) and inhibitory (K-to-Q) mutations in Stu2. As *STU2* is an essential gene, viability is supported using a *URA3* plasmid containing *STU2* (pRM10693). Yeast strains also contained *LEU2* plasmids with WT-*STU2* (yRM12337), *LEU2+* empty vector (yRM12338), Stu2-K252R (yRM12340), Stu2-K469R (12341), Stu2-K870R (12342), Stu2-2KR (yRM12343), Stu2-3KR (yRM12344), Stu2-K252Q (yRM12345), Stu2-K469Q (yRM12346), Stu2-K870Q (yRM12347), Stu2-2KQ (yRM12348), and Stu2-3KQ (yRM12349). Cells were transferred to plates lacking leucine and uracil and grown at 30 °C (B) or leucine deficient 5-FOA plates and incubated at (C) 23 °C, (D) 30 °C, and (E) 37 °C. (F,G) Following 5-FOA selection, WCEs from yeast expressing untagged Stu2 (yRM12359), Stu2-HA (yRM12358), K252R (yRM12360), K469R (12361), K870R (12362), 2KR (yRM12363), 3KR (yRM12364), K252Q (yRM12365), K469Q (yRM12366), K870Q (yRM12367), 2KQ (yRM12368), and 3KQ (yRM12369) were western blotted with anti-HA to detect relative abundances of Stu2 acetylation state mimetic mutants to the WT control.

As Stu2 is essential for yeast survival, we tested whether these alleles could support life as the sole copy of Stu2 using a 5-FOA plasmid shuffle assay (Figure 4C). As shown in Figure 4D-G, they did. All of the single, double and triple mutants support life at 23, 30, and 37 °C and displayed little or no observable differences in growth assays on a plate. These data suggest that these acetylation site mutations do not disrupt the essential function of Stu2.

### Lysine 252, 469, and 870 account for a fraction of Stu2 acetylation

To determine whether we had identified all the Stu2 acetylation sites, Stu2 was immunoprecipitated from strains expressing wild type or the 3KR and 3KQ mutants. Analysis of the precipitate with anti-HA and anti-AcK, revealed only a moderate reduction in anti-acetyl K reactivity of the Stu2 band for 3KR and only a slight reduction for the 3KQ mutant (Figure 2C and Figure S1D-F). Because acetylation reactivity was not abolished in the 3KR and 3KQ lanes, this suggests that lysines 252, 469 and 870 do not account for all of Stu2’s acetylation. We anticipate that additional acetylation site(s) are present on Stu2 and additional mass spectrometry work is in progress to identify these.

### Acetylation mutants do not display a mating defect

To begin to assess the function of the three lysines, we initially assayed three phenotypes, mating, cell cycle progression, and nuclear positioning. The fusion of nuclei during the mating process, known as karyogamy, requires intact microtubule function (Kurihara et al., 1994). To ascertain whether the acetylation of Stu2 impacts karyogamy, we employed a bilateral karyogamy mating assay in which both the *MAT*a and *MAT*α partners contained the same *stu2* mutation. In comparison to *kar9*Δ, which is known to exhibit a moderate karyogamy defect (Kurihara et al., 1994) (Miller & Rose, 1998), we observed little or no mating defect for K or Q substitutions at K252, K469 or K870 in Stu2 (Figure 5A).

**Figure 5.**
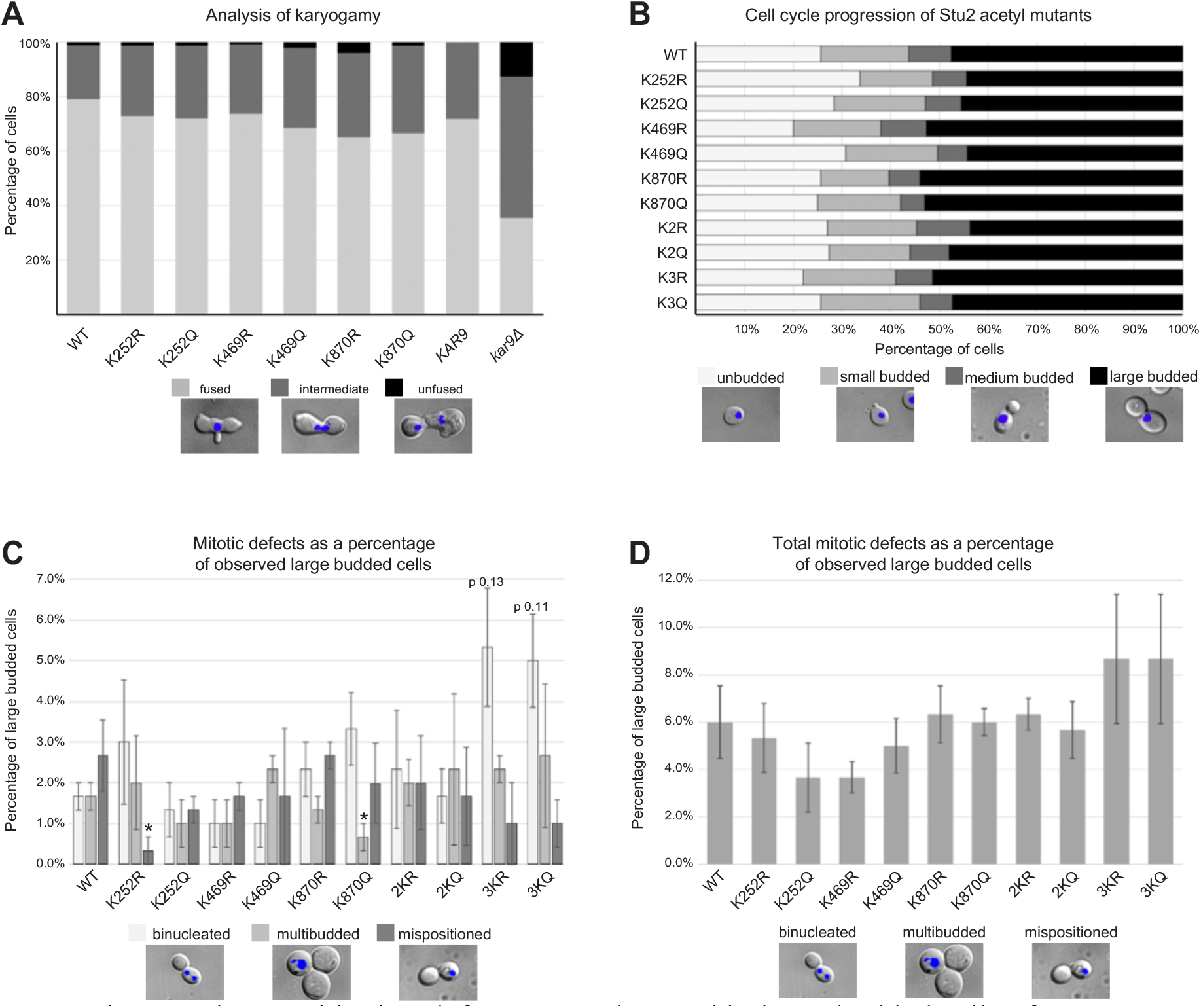
Minor nuclear positioning defects were observed in large-budded cells of yeast expressing the 3KR and 3KQ Stu2 acetyl mutations. (A) *MAT*a or *MAT*α mating type yeast with plasmids expressing *STU2* (yRM11379 and yRM11368), Stu2-K252R (yRM11930 and yRM11892), Stu2-K252Q (yRM11934 and yRM11898), Stu2-K469R (yRM11375 and yRM11362), Stu2-K469Q (yRM11377 and yRM11366), Stu2-K870R (yRM11938 and yRM11912), Stu2-K870Q (yRM11940 and yRM11922) in a *stu2Δ* background were tested in a karyogamy assay as described in (Miller & Rose, 1998). In addition, yeast containing a genomic copy of *KAR9* (yRM299 and yRM301) or a genomic deletion of *KAR9* (yRM393 and yRM396) were assayed to represent moderate karyogamy defects. (B) Logarithmically growing yeast with plasmids expressing *STU2* (yRM12358), K252R (yRM12360), K469R (12361), K870R (12362), 2KR (yRM12363), 3KR (yRM12364), K252Q (yRM12365), K469Q (yRM12366), K870Q (yRM12367), 2KQ (yRM12368), and 3KQ (yRM12369) in a *stu2Δ* background were evaluated for the percentage of unbudded, small budded, medium budded and large budded cells in the population. (C) Large-budded cells of yeast were DAPI stained and further examined for binucleate, multi-budded, and cell polarity defects. Bar graphs displayed as mean ± SEM derived from 3 counts of 100 large budded cells; * p < 0.1. 3KR and 3KQ exhibiting binucleation defects were close, but did not pass t-testing thresholds for marginal significance with p values of 0.13 and 0.11 respectively. (D) The aggregate of mitotic defects across Stu2 acetylation mutants are graphed for comparison to a WT control.

Next, we asked whether acetylation altered the cell-cycle progression by analyzing the distribution of unbudded, small budded, medium budded, or large budded cells in an actively growing culture. As shown in Figure 5B, little or no observable defect was seen. However, we observed an increase in aberrantly positioned nuclei in large-budded cells. In these cells, there was an increase in the number of mother cells containing two nuclei, referred to as binucleate cells (Miller & Rose, 1998). There was also an increase in the number of cells that appeared multi-budded (Figure 5C). Furthermore, microscopy of mispositioned nuclei revealed that while binucleated populations were potentially elevated in the 3KR and 3KQ mutants (Figure 5C), there was little to no statistical significance between any of the mutants and the WT control (p=0.13 and p=0.11for 3KR and 3KQ respectively), even when the three phenotypes are viewed in aggregate (Figure 5D).

### A subset of Stu2 acetylation mutations confer benomyl resistance

Stu2 promotes microtubule polymerization (Huffaker et al., 1988) (Podolski et al., 2014). If these lysine mutations were affecting this function of Stu2, we theorized that they might be more sensitive to drugs that destabilize microtubules. We first tested the effect of benomyl on the acetyl inhibitory mutants, (K-to-R). As shown in Figure 6, the K252R, the K469R, and their corresponding double mutant in the TOG domains, 2KR, grew similarly or only slightly better than wild type. Surprisingly, the K870R and the 3KR mutant grew much better than wild type. As the triple 3KR mutant also displays benomyl resistance like the single K870R mutant, it suggests that acetylation at K870R is intragenetically epistatic to acetylation in the TOG domains. In other words, the K870R phenotype predominates over that of the 2KR in this assay. Since these mutants were assayed for benomyl resistance in haploid *stu2Δ*, with the mutations present on a YCP plasmid as the cells only source of Stu2p, it also suggests the likelihood that the K870R mutation stabilizes microtubules.

**Figure 6.**
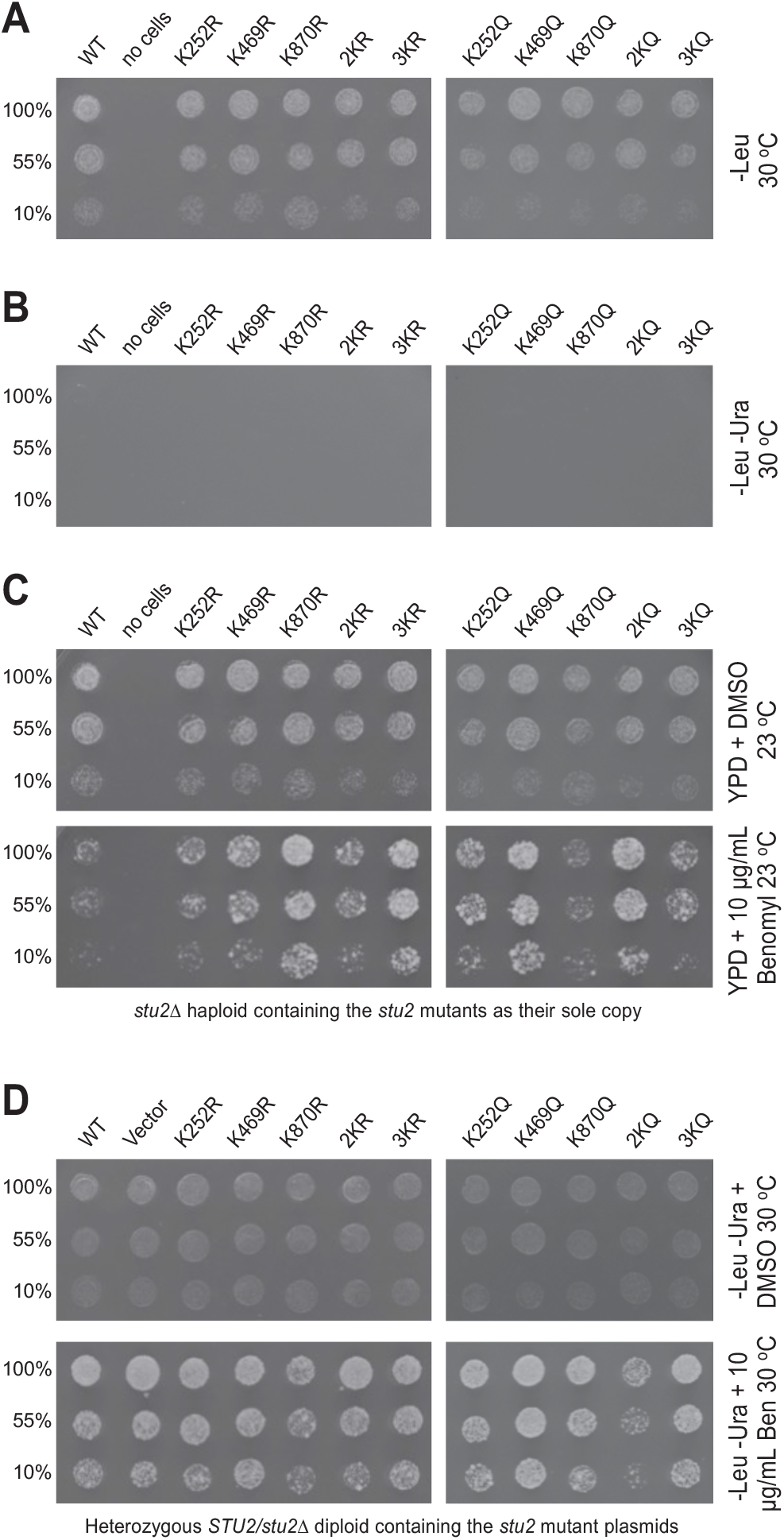
Some acetylation mutants display resistance to the microtubule destabilizing drug, benomyl. Yeast containing *LEU2* plasmids with *STU2* (yRM12358), K252R (yRM12360), K469R (yRM12361), K870R (yRM12362), 2KR (yRM12363), 3KR (yRM12364), K252Q (yRM12365), K469Q (yRM12366), K870Q (yRM12367), 2KQ (yRM12368), and 3KQ (yRM12369) were transferred to (A) plates lacking leucine and (B) plates lacking leucine and uracil to confirm removal of wild-type *STU2 URA3* plasmids. (C) Cells were also transferred to YPD containing DMSO or DMSO + 10 μg/mL benomyl and grown at 23 °C. (D) Yeast containing *LEU2*+ plasmids with *STU2* (yRM12233), *LEU2+* empty vector (yRM12234), Stu2- K252R (yRM12241), Stu2-K469R (yRM12239), Stu2-K870R (yRM12252), Stu2-2KR (yRM12247), Stu2-3KR (yRM12250), Stu2-K252Q (yRM12242), Stu2-K469Q (yRM12240), Stu2-K870Q (yRM12251), Stu2-2KQ (yRM12248), and Stu2-3KQ (yRM12249) in a heterozygous delete *STU2*/*stu2-Δ1*::*HIS3* background at the genomic locus were transferred to –leu –ura plates containing DMSO or DMSO + 10 μg/mL benomyl and grown at 30 °C.

We next tested whether the acetyl-mimetic mutations would affect Stu2 sensitivity to benomyl. Both the K252Q and K469Q single mutations, as well as the 2KQ double mutation, conferred resistance to benomyl (Figure 6C). In contrast to the K870R mutation, the K870Q mutation did not display resistance to benomyl. To ascertain which phenotype is intragenetically epistatic, we again compared the 2KQ double TOG mutation to the 3KQ mutant. In this analysis, the 3KQ mutant displayed a similar phenotype to the K870Q single mutant with respect to its level of benomyl resistance, suggesting that the K870Q phenotype is epistatic to that of the double Q mutant, K2Q. Notably, the phenotype displayed by both the R and Q configurations at position 870 overrides the phenotype of the corresponding double mutant in the TOG domains. These phenotypes were also visible at 30 °C (Figure S2). Together, these data suggest that acetylation in both the TOG domain at K469 and MAP binding domain at K870 regulate the ability of Stu2 to stabilize microtubules. It is noteworthy that for benomyl resistance, acetylation at these two domains appear to function in opposition to each other. It remains to be determined whether the cycling of acetyl moieties on-and-off of Stu2 plays a role in microtubule stability.

As Stu2 is known to function as a dimer, we next sought to determine whether the benomyl- resistance phenotypes are the result of either dominant or recessive mutations. For this, the benomyl phenotypes were assayed in a heterozygous *STU2*+/*stu2Δ* diploid strain containing the Stu2 mutant plasmids. As shown in Figure 6D, all of the mutants grew at rates comparable to wild type, except 2KQ, which showed a moderate inhibition of growth. This suggests that the 2KQ mutant is partially dominant. Moreover, several of the benomyl phenotypes differed between the *stu2Δ* and strains containing a wild-type genomic copy of *STU2* (compare panel 6C to 6D). This implies that symmetrical acetylation is important between the two halves of a Stu2p dimer.

### Impaired Stu2 acetylation leads to chromosome instability

As Stu2 plays an important role in linking microtubules to the kinetochore, we asked whether the acetylation state of Stu2 can compromise sister chromatid segregation. To test this, we employed the acetylation mimetic and acetylation preventative mutants in an artificial chromosome loss assay using a quantitative 5-FOA counter-selection, as depicted in Figure 7A. This standard assay employed yeast containing a 125 kb artificial chromosome that was marked with *URA3* in a *stu2-Δ1*::*HIS3* background, as described in Materials and Methods (Shero et al., 1991). Orotidine-5’-phosphate (OMP) decarboxylase encoded by the *URA3* gene converts 5- FOA into 5-Fluorouracil, a toxic compound. Yeast can only grow on media containing 5-FOA in the absence of the *URA3*-marked artificial chromosome, providing a quantitative assessment of chromosome transmission fidelity.

**Figure 7.**
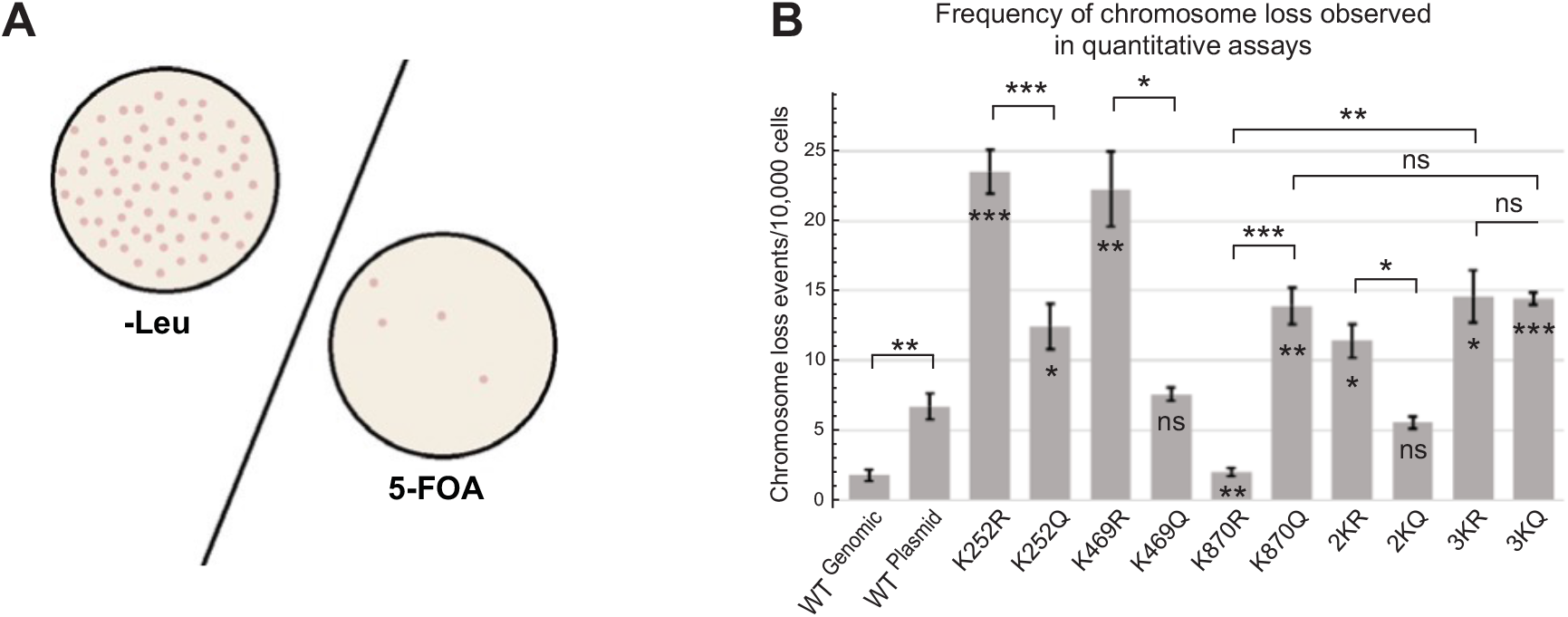
Manipulation of Stu2 acetylation sites alters chromosome transmission fidelity. Yeast strains for this assay contained a 125 kb artificial chromosome that expressed the *URA3* gene, in addition to plasmids expressing WT-Stu2 (yRM12253), K252R (yRM12261), K252Q (yRM12262), K469R (yRM12301), K469Q (yRM12260), K870R (yRM12271), K870Q (yRM12270), 2KR (yRM12266), 2KQ (yRM12267), 3KR (yRM12269), and 3KQ (yRM12268) in a *stu2*-Δ1::*HIS3* background. We also evaluated the chromosome loss rate when the wild type *STU2* was present at its genomic locus with an empty *LEU2* vector (yRM12300). (A) To quantify the frequency at which for the artificial chromosome was lost in Stu2 mutants, the cells were transferred to plates containing 5-FOA or serially diluted and transferred to -leu synthetically defined plates. Because 5-FOA is toxic to yeast possessing the *URA3* gene, only colonies that have lost the 125 kb artificial chromosome should grow on 5-FOA. Colonies growing on leucine deficient plates reflect the total number of cells used in the experiment. (B) Quantitative rates of chromosome transmission fidelity reflect the total number of colonies growing on 5-FOA media relative to colonies growing on -Leu synthetic deficient media following serial dilutions. Quantitative chromosomal loss data displayed as mean ± SEM of quadruplicate plates; * p < 0.05, ** p < 0.01, *** p < 0.005 (raw quantitative data are available in supplementary file 1).

As shown in Figure 7B, the single acetyl-preventative mutants K252R and K469R showed the largest decrease in chromosome transmission fidelity. In comparison, the acetyl- mimetic mutation K252Q showed a moderate decrease in chromosome transmission fidelity, whereas K469Q showed no significant change when compared to the plasmid-based WT-Stu2 control. The acetylated-state mimetic K870Q caused a moderate decrease of chromosome fidelity, similar to the K252Q mutant. In contrast, the K870R mutation improved chromosome transmission fidelity significantly, matching the rate observed when the wild-type *STU2* was present at its genomic loci. Double TOG domain mutants 2KR and 2KQ displayed mild phenotypes relative to the single corresponding TOG domain mutants. Lastly, when all three acetylation sites were mutated to arginine or glutamine, moderate decreases in chromosome transmission fidelity were observed. Further, similar trends were observed when the assay was performed qualitatively using red sectoring of colonies for the mutants (Figure S3). Combined, these results suggests that Stu2 acetylation plays an important role in controlling chromosome loss. We speculate that this may be due to regulation of Stu2’s protein-protein interactions important for microtubule attachment at the kinetochore.

### Stu2 acetylation regulates interactions with γ-tubulin

In addition to its localization at kinetochores and at MT plus ends, Stu2 also functions at the SPB (Wang & Huffaker, 1997). Stu2 is known to interact with the γ-TuSC of which γ-tubulin is a part (Geissler et al., 1996) (Knop et al., 1999) (Knop et al., 1997) (Vinh et al., 2002) (Usui et al., 2003). Three recent studies provide direct evidence for Stu2/XMAP215 roles in microtubule nucleation at the SPB/MTOC. In yeast, Stu2 promotes γ-TuSC mediated MT nucleation (Gunzelmann et al., 2018) (Brilot et al., 2021) and in *X. laevis*, XMAP215 played a similar role at the more elaborate γ-TuRC (Thawani et al., 2018).

Because of the similarity of γ-tubulin with the other classes of tubulin, we sought to gain insight into the interactions of the Stu2 mutants with its tubulin binding partners. For this, we first aligned β- and γ-tubulins from seven eukaryotes across evolution (Figure 8A,B). It has long been recognized that these tubulins are highly conserved from fungi to humans. Previous crystallography work from the Rice laboratory elucidated the position of residues E158-D161 within β-tubulin (Ayaz et al., 2014), denoted here as the EFPD loop. Their modeling predicts that this loop interacts with Stu2’s TOG2 near K469. Based on this work, we then identified the analogous loop within γ-tubulin as residues R166-K169, denoted here as the RYPK loop. We hypothesize that this loop in γ-tubulin also docks with Stu2.

**Figure 8.**
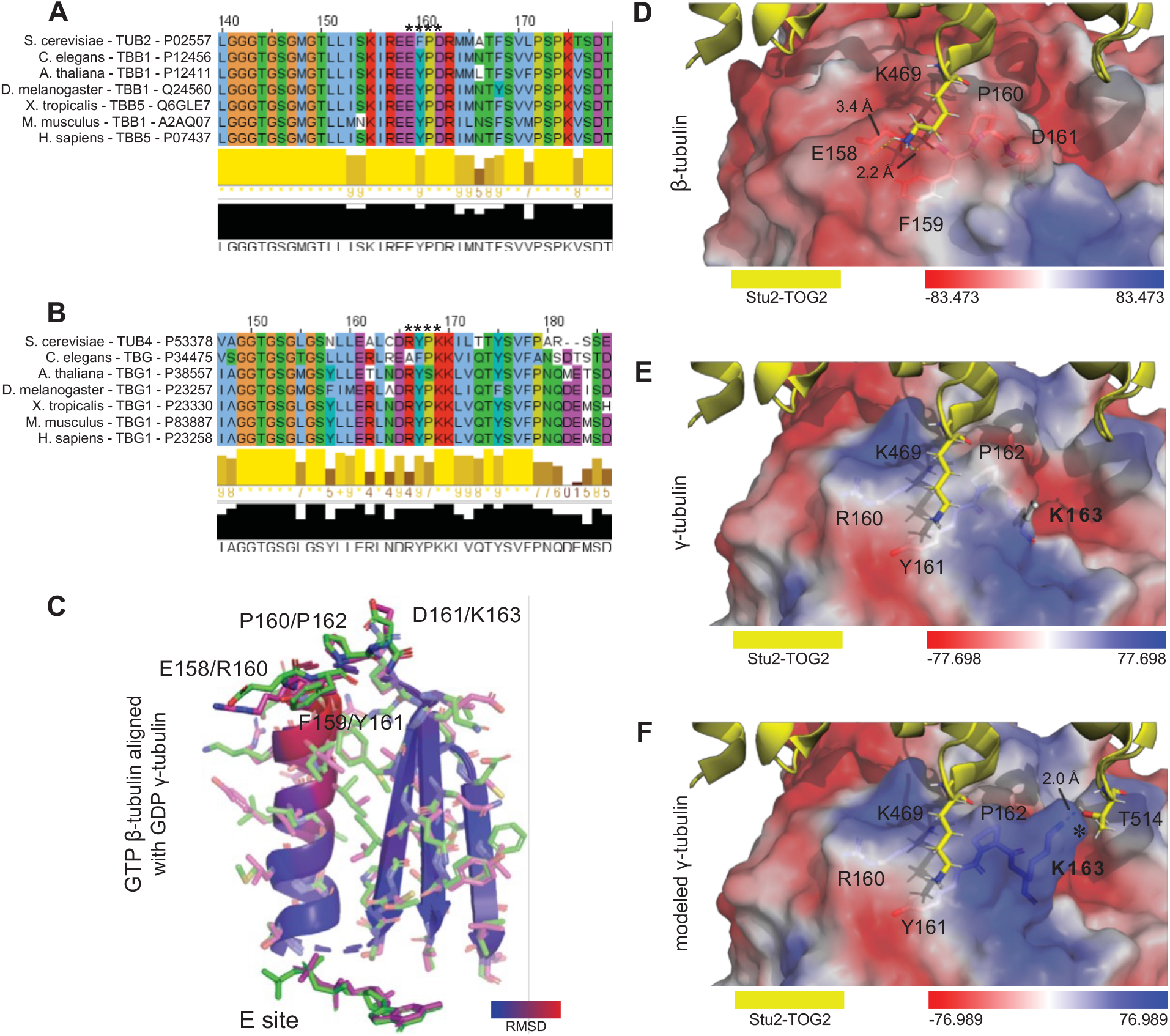
K469 is predicted to interact with the evolutionarily conserved EFPD and RYPK loops of β-and γ-tubulin. We evaluated evolutionary conservation of eukaryotic (A) β-tubulin EFPD and (B) γ-tubulin RYPK loops that are predicted to be responsible for tubulin interactions with Stu2 K469. (C) Structural alignments show similarities between GTP bound β-tubulin EFPD (pdb:4U3J) and GDP bound γ-tubulin RYPK (pdb:3CB2) loops. Electrostatic charge maps illustrate significant differences between the (D) electronegative β-tubulin EFPD loop (pdb:4U3J) and the (E) electropositive GCP2/Spc97 bound γ-tubulin RYPK loop in a closed confirmation γ-TuSC (pdb:5FLZ). To portray more accurately the positively charged surface of γ-TuSC embedded γ-tubulin, an electrostatic surface map (F) was generated following γ-tubulin K163 side chain reconstruction. The K163 sidechain of γ-tubulin is in position to form an additional salt bridge with T514 found in the Stu2 TOG2 domain (panel F marked with *).

To evaluate conservation of interactions in these regions between tubulins, we aligned the crystal structures of γ-tubulin (pdb:3CB2) with β-tubulin (pdb:4U3J) using PyMOL (Figure 8C) (Ayaz et al., 2014) (Rice et al., 2008). This alignment illustrates a high degree of similarity between the two loops in β- and γ- tubulins for the peptide backbone and sidechains. Notably however, the EFPD (β-tubulin) and RYPK (γ-tubulin) loops display a significant difference in their ionic charge. In the case of β-tubulins, the residues of the EFPD loop are acidic with a net charge -2 and in the case of γ-tubulins the residues of the RYPK loop are basic with net charge +2. As described below, we theorized that charge modulation at K469 influences interaction with these loops.

While evaluating the interactions of K469 with β- and γ-tubulins, we observed that the side chain of K469 was not solved in the crystal structure. To gain insight into this region, we used PyMOL to reconstruct the K469 side chain and screened for optimal rotamer confirmations. As shown in Figure S4F and G, this analysis revealed that Stu2’s K469 is capable of forming salt bridges with β-tubulin residues E158 and aspartate D161. To visualize these charge arrangements, we also compared the position of the reconstructed K469 side chain with electrostatic surface maps of β- and γ-tubulins. As shown in Figure 8C and D, the surface charges of the two tubulins differed significantly in the region surrounding the loops. The β- tubulin is negatively charged, and the γ-tubulin is positively charged.

The surface map of β-tubulin with TOG2 in this region is well-characterized, based on crystallography data (Ayaz et al., 2014). However, during our analysis of the surface charges present on the γ-tubulin RYPK loop, we noticed the orientation of the side chain of γ-tubulin’s K163 remained unsolved (Greenberg et al., 2016) (Figure 8D). To fill this lacuna, we further modeled the γ-tubulin K163 side chain using PyMOL to reconstruct possible side chain data. As seen the model depicted in Figure 8E, the positive charge of arginine 160 and lysine 163 would likely repel the positive side chain of K469 found in TOG2.

If Stu2 interacts with both β- and γ-tubulin through this motif, then our model predicts that acetylation at K469 will affect each of the interactions differently because the charge of the EFPD and RYPK loops are different. In this model for γ-tubulin, Stu2 interaction should be promoted by the acetylation of K469 (imitated by Q). This would be due to reduced ionic repulsion between the acetyl-lysine of Stu2 and the positively charged residues within the RYPK loops of γ-tubulin. However for β-tubulin, the non-acetylated form of lysine (mimicked by R) should be preferred over an acetylated lysine.

To test this prediction, we assayed the interaction of Stu2 and the tubulins with a co-immunoprecipitation assay. We precipitated wild-type Stu2, as well as each of the K-to-R and K-to-Q mutants, and then assayed for γ-tubulin in the precipitates. Whereas the single K-to-R mutant K469R (as well as the K252R and K870R single mutants) consistently showed little or no reduction in the levels of γ-tubulin co-immunoprecipitated (Figure 9A), the 3KR displayed a reproducible reduction compared to wild type. In contrast, the K469Q mutation, as well as the 2KQ and 3KQ mutants, all displayed elevated enrichment of γ-tubulin. The strongest enrichment in γ-tubulin binding is seen in the 3KQ mutant (Figure 9A). The strongest decrease in γ-tubulin binding is displayed by the 3KR mutant. Combined, these findings suggest that acetylation regulates the interaction between Stu2 and γ-tubulin.

**Figure 9.**
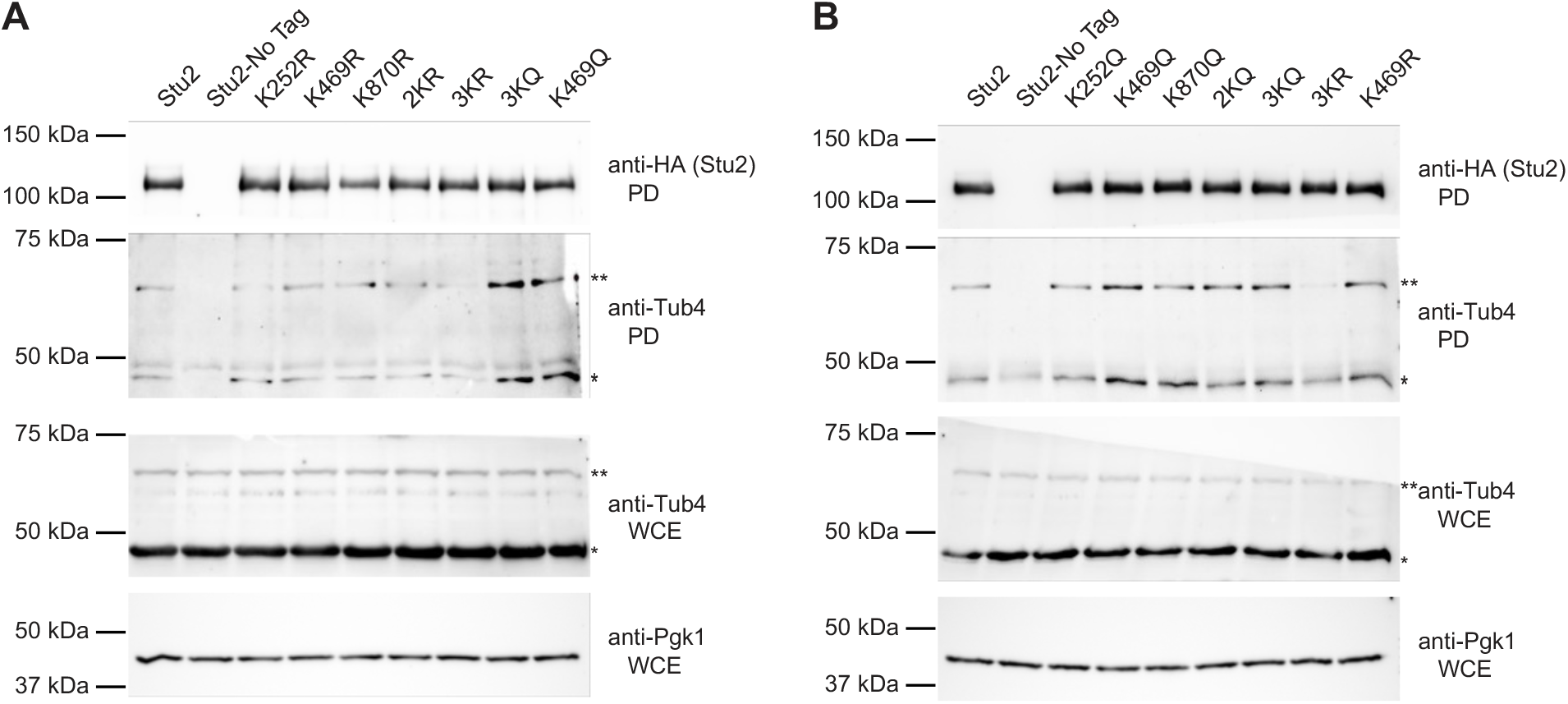
The acetyl-mimetic K469Q and 3KQ mutants promote Stu2 interactions with γ- tubulin. (A,B) Yeast expressing Stu2-HA (yRM12358), untagged Stu2 (yRM12359) K252R (yRM12360), K469R (12361), K870R (12362), 2KR (yRM12363), 3KR (yRM12364), K252Q (yRM12365), K469Q (yRM12366), K870Q (yRM12367), 2KQ (yRM12368), and 3KQ (yRM12369) were immunoprecipitated in PBS buffer and western blotted for HA tagged Stu2. To detect co-precipitation of γ-tubulin, pull-downs were also probed with anti-Tub4. To evaluate γ-tubulin abundance and confirm uniform loading of protein, WCEs were western blotted for γ-tubulin and Pgk1.

In these blots, it is evident that an additional higher-molecular weight band of γ-tubulin that runs at 62.5kDa co-precipitates with Stu2. As γ-tubulin has been reported previously to be ubiquitinated in mammalian cell lines HEC293T cells (Yin et al., 2021) (Starita et al., 2004) (Sankaran et al., 2005) (Sankaran et al., 2007), we speculate that this may be the molecular basis of the shift seen here. Notably, it appears that the modified form of γ-tubulin is enhanced in the interactions with Stu2 (Figure 9A,B, compare γ-tubulin in the pulldowns to the WCEs).

The model also predicts that the positive charge on K469 is important for Stu2 interactions with β-tubulin. To test this prediction, we assayed the Stu2 precipitates for the presence of β-tubulin, using its α-tubulin partner as a surrogate. As seen in Figure 10, slightly less α-tubulin was present in the immune-precipitate of K469Q than wild type. Although the reduction was only moderate, this reduction was reproducibly consistent. K469R interacted with Stu2 at a level equivalent to wild-type Stu2. This suggests that acetylation of K469 regulates interactions with both β-tubulin and γ-tubulin.

**Figure 10.**
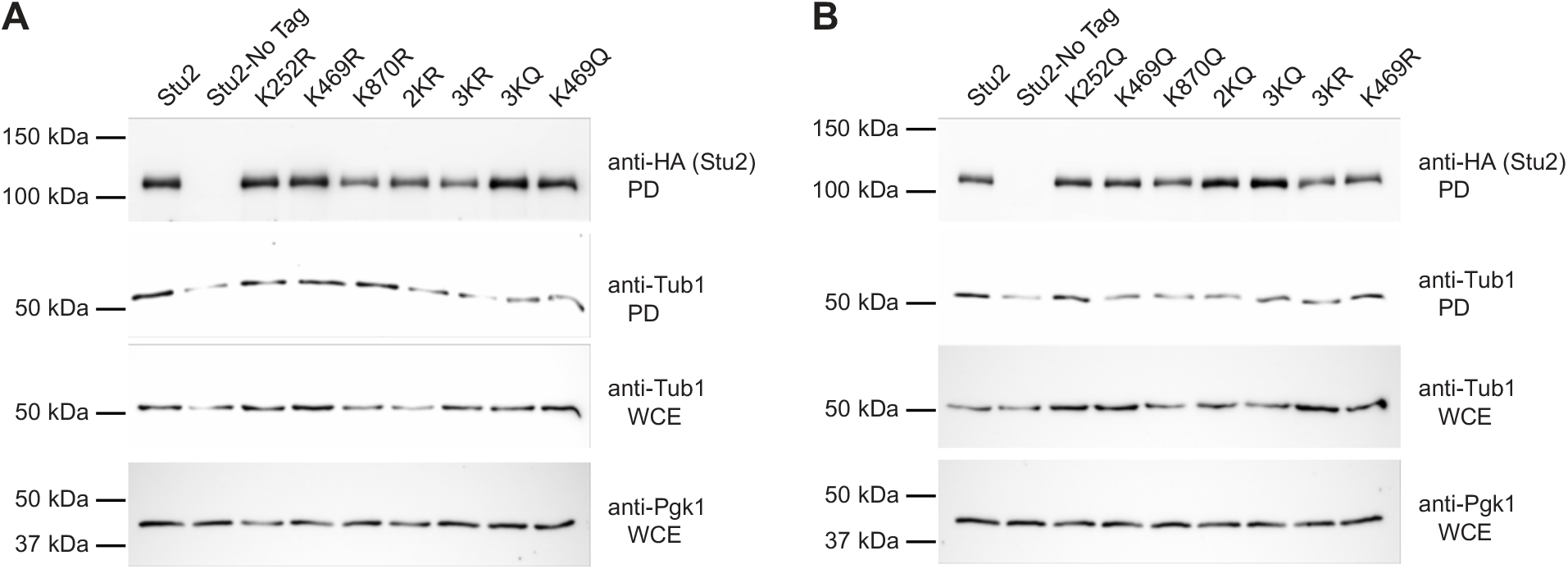
The interactions of Stu2 acetyl-mimetic mutants with αβ-tubulin. (A,B) The yeast described in Figure 9 were also used for Stu2 immunoprecipitations in PEM buffer and western blotted for HA tagged Stu2. To detect co-precipitation of αβ-tubulin, pull-downs were also probed with anti-Tub1. To evaluate α-tubulin abundance and confirm uniform loading of protein, WCEs were western blotted for α-tubulin and Pgk1.

The model also predicts that the positively charged K469 is important for Stu2 interactions with β-tubulin. To test this prediction, we assayed the Stu2 precipitates for the presence of β-tubulin, using its α-tubulin partner as a surrogate. As seen in Figure 10, slightly less α-tubulin was present in the immune-precipitate of K469Q than wild type. Although the reduction was only moderate, this reduction was reproducibly consistent. K469R interacted with Stu2 at a level equivalent to wild-type Stu2. This suggests that acetylation of K469 regulates interactions with both β-tubulin and γ-tubulin.

## DISCUSSION

This work represents the first functional characterization of acetylation for any Stu2/XMAP215 family member. Our report details how acetylation regulates multiple functions of the Stu2 protein. Using a collection of acetyl-inhibitory (K-to-R) and acetyl-mimetic (K-to-Q) mutants, we show that acetylation of three lysines within Stu2 at K252, K469, and K870 is important for microtubule stability, chromosome segregation, and affinity for its γ-tubulin binding partner. Collectively, these findings suggest that multiple activities of the Stu2 protein are coordinated by acetylation, including those at the kinetochore and SPB, and these are summarized in Figure 11A.

**Figure 11.**
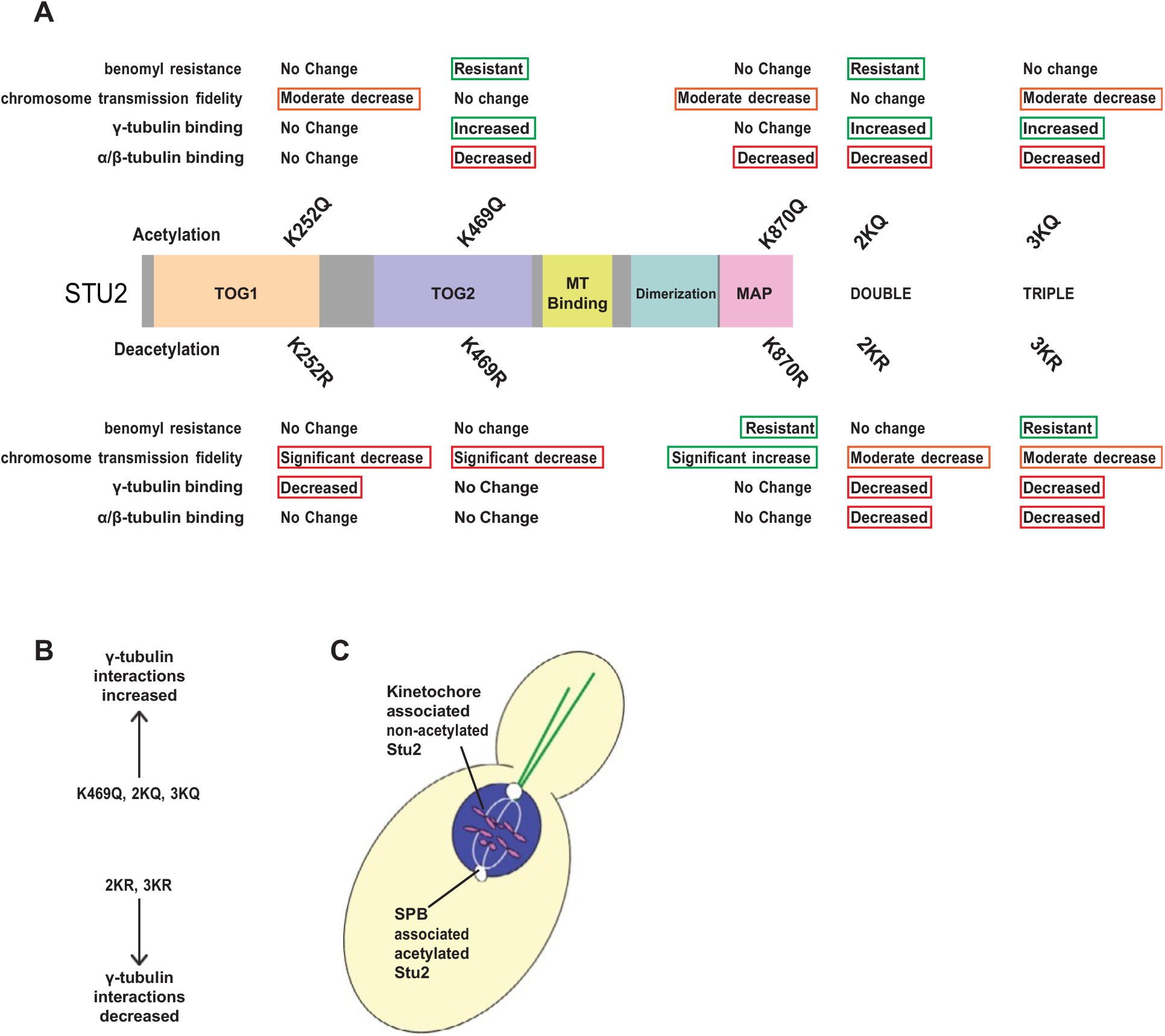
Acetylation regulates multiple functions of Stu2. (A) Experimental results are summarized for the Stu2 mutants in assays for benomyl sensitivity, chromosome transmission fidelity, and interactions with β- and γ-tubulin. (B) Depicts a summary of co-immunoprecipitated γ-tubulin by key Stu2 mutants compared to the Stu2 wild type controls. (C) A working model of our acetylation hypothesis is shown. This model predicts that de- acetylation of Stu2 promotes its interaction with kinetochores and that Stu2 acetylation promotes interaction with the γ-tubulin complex.

Several of the mutants display improved Stu2 function. (eg. K870R for chromosome transmission fidelity; K469Q and K870R for benomyl resistance; and 3KQ and K469Q for interactions with γ-tubulin). This suggests that these effects are not simply the result of compromised protein function, but rather they represent modulation of Stu2 activity based on regulation of critical ionic charges.

Our analysis of these lysines suggest that several functions of Stu2p can be separated based on their acetylation status. This was particularly true for acetylation at K469 and K870. The influential nature of K870 acetylation was especially evident in the benomyl sensitivity assays. At this residue, the Q and R states produced distinctly different phonotypes. Regardless of its acetylation status, the K870 mutation suppresses the benomyl phenotype produced by corresponding double mutation in the TOG domain (the 2K). For example, the non-acetylatable K870R produces robust benomyl resistance, whereas the 2KR mutant does not. When the K870R is added to the 2KR to produce the triple (3KR) mutants, the strain is again benomyl resistant. A similar pattern is seen for the acetyl-mimetic K870Q (Figure 6C). Thus, this suggests that the acetylation state of K870 effectively overrides the information contained within the TOG-domain acetylation sites for stabilization of microtubules.

Moreover, the pseudo-acetylation of the three lysines all trend toward increasing interactions with γ-tubulin. Notably, the pseudo-acetylation mutants K469Q and 3KQ have the largest increases (Figure 9). Although the single K to R mutants display little to no reduction in γ-tubulin binding, the 3KR mutant displays a significant reduction in γ-tubulin binding.

Our data also shows that Stu2 acetylation regulates chromosome transmission fidelity, both positively and negatively. Acetylation at K252 and K469 found in the TOG1 and TOG2 domains respectively were both important for chromosome segregation, since mutants unable to undergo acetylation at these sites displayed decreased chromosome transmission fidelity. In contrast, the non-acetylatable surrogate, K870R, displayed significantly better chromosome transmission rates than the constitutively acetylated surrogate, K870Q (Figure 7B).

In addition to the Ndc80 complex interacting through the c-terminal MAP domain, Stu2 also interacts with Bik1 and Bim1 through this domain (Wolyniak et al., 2006). These MAPs serve roles in tracking and stabilizing microtubule plus ends (Carvalho et al., 2004) (Wolyniak et al., 2006) (Blake-Hodek et al., 2010) reviewed in (Akhmanova & Steinmetz, 2008). Because lysine 870 is located within this domain, we speculate that this lysine may mediate interactions with these MAP proteins.

Recent novel findings describe how Stu2 functions at the kinetochore as a mechanosensor to promote correct bilateral anchoring of sister-chromatids to opposing spindle pole buddies in yeast undergoing mitosis (Miller et al., 2016) (Miller et al., 2019) (Humphrey et al., 2018).

However, future work will be needed to determine whether acetylation regulates the mechano- sensing ability of Stu2. Work from Matthew Miller’s lab also demonstrated that K870 resides in a patch of amino acids critical for Stu2 interactions with the Ndc80 complex (Zahm et al., 2021). In addition to our alignments of XMAP215 family members, recent work by Zahm *et al*. (2021) showed that several species including the yeast *K. marxisus*, zebra fish *D. rerio*, and the bird *G. gallus* each possess a region from L869-K877 that is important for association with the Ndc80 complex. Our work showing that K870 is highly conserved agrees with their alignments of the region (see Figure 1C and Figure 4F of (Zahm et al., 2021)), and raises the possibility that acetylation at K870 may likewise be highly conserved. Additionally, Zahm et al showed that Stu2 K870 forms salt bridges with Spc24, and that Stu2 R872 also forms salt bridges with adjacent D651 and S654 residues of Ndc80. This indicates that the three amino acids K870 A871 R872 likely play a role in Stu2, Ndc80, and Spc24 complex integrity. Lastly, in three of the fungi and three of the eukaryotic strains, we note that the amino acids in the K870 A871 R872 motif are flipped to R870 A871 K872. We speculate that in some of these systems a putative acetylation at a K872 may target that face of the Stu2 alpha helix to impair salt bridges with Ndc80 rather than those of Spc24. Nevertheless, these findings make the prediction that acetylation at K870 will regulate these Ndc80-complex dependent interactions.

### Modeling the role of acetylation in regulating Stu2’s interaction with the γ-TuSC

Our conclusion that acetylation of Stu2 at K469 regulates its interactions with different tubulins is also informed by modeling of these interactions using previously published structures (Ayaz et al., 2014) (Greenberg et al., 2016). In examining these structures, we noticed that Stu2’s K469 interacts with tubulin at a specific set of tubulin residues, termed EFPD and RYPK loops of β- and γ-tubulin respectively. These loops forming a binding interface with Stu2’s K469 possess significantly different charges. We hypothesize that the charge negation that acetylation imparts drives Stu2-interaction with γ-tubulin. Further, the labile nature of acetylation may explain previous difficulties in observing Stu2 interactions with γ-tubulin. It is possible that acetylation-mediated interactions with γ-tubulin are part of the mechanism by which Stu2 is incorporated into the γ-tubulin small complex (γ-TuSC), in addition to its interactions with Spc72. We are currently investigating the interactomes of the Stu2 acetyl- mutants to test this idea.

To frame these interactions in the context of MT nucleating complexes, we modeled Stu2 interactions with γ-tubulin using PyMol. For this, we juxtaposed the structure of the β- tubulin/TOG complex derived from x-ray crystallography coordinates found in pdb:4U3J (Ayaz et al., 2014) with the structures of γ-TuSC that were generated from the cryo-EM of Greenberg *et al.,* (2016). We did this for both their open and closed conformations, described by the coordinates in pdb:5FM1 and pdb:5FLZ, respectively. Our model depicting these structures is shown in Figure 12A and B. We used the root-mean-square deviation to color code positional conservation of the structures of β- and γ-tubulin, as described in materials and methods (Ayaz et al., 2014) (Greenberg et al., 2016). For these tubulins, the degree of structural conservation is shown on a scale ranging from blue to red. Similar structure is depicted in blue and non-conserved structure is depicted in red. This modeling indicates two things. First, the binding interface between the Stu2 TOG2 domain and γ-tubulin resembles that of β-tubulin. Second, Spc97 and Spc98 are positioned to form contacts with the binding interface of TOG2 that would normally be occupied by the α-tubulin found in an αβ-tubulin heterodimer.

**Figure 12.**
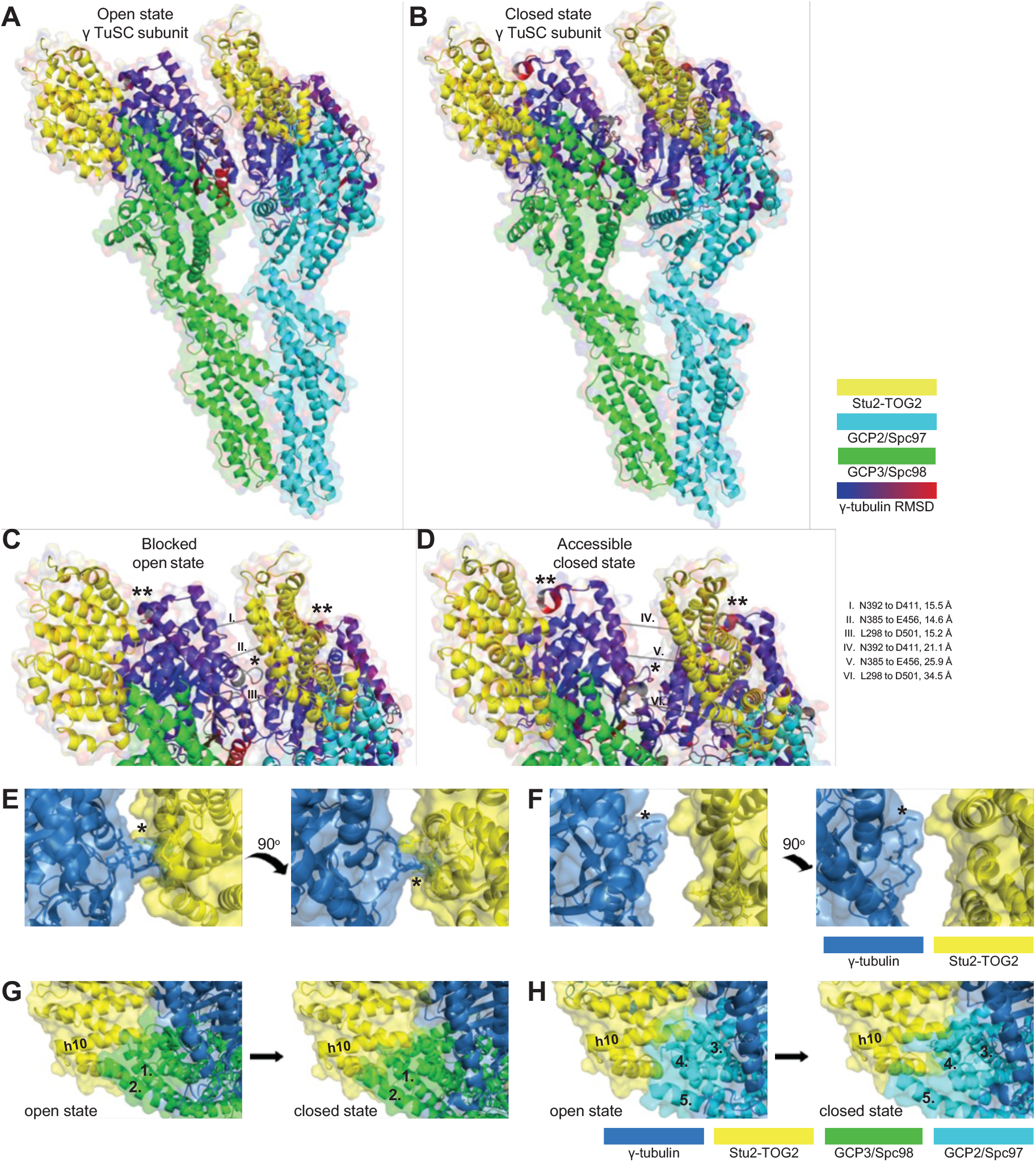
A composite model derived from cryo-EM and x-ray crystallography structures predicts that Stu2’s TOG interactions may be regulated by the open/closed conformational states of the γ-TuSC. A TOG bound γ-TuSC subunit is depicted in the (A) open state, adapted from pdb:5FM1 and pdb:4U3J, and (B) closed state, adapted from pdb:5FLZ and pdb:4U3J. A close-up view of the γ-TuSC complex in the (C) open state, and (D) closed state reveals SPB state dependent TOG domain accessibility. Distances between secondary amines of peptide backbones are reported in angstroms for the listed amino acids. A structured loop from the adjacent γ-tubulin projects into the space theoretically occupied by the Stu2 TOG2 domain in the open (E) but not the closed (F) confirmation γ-TuSC structure (marked with * in panels C-E). Stick structures are shown for γ-tubulin residues S307 through P313 and Stu2 TOG2 residues E456, L459, K460, R497, Y498, and E502. Relaxation of steric hindrance between TOG2 α-helix 10 of Stu2 and two α-helices of Spc98, corresponding to 1. aa Q756-L779 and 2. aa S798- D833, occurs as γ-TuSC undergo open to closed conformational changes (G). Relaxation of TOG2 α-helix 10 of Stu2 and three α-helices of Spc98, corresponding to 3. aa S575-K608, 4. aa L692-S712, and 5. aa E754-E785, similarly occurs as γ-TuSC undergo open to closed conformational changes (H). Additional contacts are predicted between γ-tubulin in closed confirmation γ-TuSC and α-helices 2 and 4 of nearby TOG2 domains as indicated with ** in panels C and D.

From this modeling, we predict that Stu2 interactions are more likely to occur with the γ- TuSCs in their closed conformation rather than in their open conformation. This concept is based upon three observations that are apparent in the model. First, in the open γ-TuSC structure, a loop within γ-tubulin (amino acids S307-N312) extends into spaces normally occupied by the TOG2 domain, which would interfere with Stu2 binding. This steric hindrance is apparent in the backbone ribbon structure of Figure 12C and indicated with an asterisk*. In the closed γ-TuSC structures, this loop is moved away from the TOG2 binding pocket (Figure 12D). For both the open and closed conformations, this loop’s accessibility is also illustrated in space filling models (Figure 12E,F).

Second, we observe in this model that alpha helix 10 of Stu2’s TOG2 domain interferes with the space occupied by multiple gamma ring proteins (GRIPs) of the γ-TuSC. Alpha helix 10 of TOG2 domains overlap with two alpha helices of Spc98 (Figure 12G) and three alpha helices of Spc97 (Figure 12H). While this is an apparent weakness with our model, we note that overall, the apparent steric hinderance is reduced as the structure goes from an open to a closed conformation.

Third in this model, a loop of γ-tubulin (comprised of amino acids D411 through S415) moves into position and forms additional binding contacts with TOG2 α-helices 2 and 4 specifically in the closed confirmation (Figure 12C,D indicated with **). In addition, the distance between neighboring γ-tubulins and TOG domains is expanded by approximately 12.1 Å when transitioning from an open to a closed conformation, theoretically improving access of the TOG2 domain to its γ-TuSC binding interfaces.

Here, we report that K252 acetylation is important for chromosomal segregation (Figure 7). However, unlike K469 and K870 for which there is a wealth of structural data and interaction mapping available (Miller et al., 2019) (Chen et al., 1998) (Slep & Vale, 2007) (Al-Bassam et al., 2007) (Al-Bassam & Chang, 2011) (Ayaz et al., 2012) (Ayaz et al., 2014) (Zahm et al., 2021) (Gunzelmann et al., 2018), little structural information is available to explain why inhibition of K252 acetylation leads to reductions in chromosome transmission fidelity. Because K252 is positioned between TOG1 and TOG2 domains, it is possible that acetylation plays a role in the “polarized unfurling” model for Stu2 as a microtubule polymerase (Nithianantham et al., 2018). Consistent with an unfurling model, our I-TASSER modeling suggests that K252 acetylation may facilitate elongation of the intra-TOG domain linker region of microtubule polymerases to reduce steric clash of bound tubulin heterodimers, since K252 acetylation is predicted to reduce the formation of salt bridges with neighboring D289 and K293 (Figure 13C,D). We speculate that acetylation may impart extra flexibility in the intra-TOG domain.

**Figure 13.**
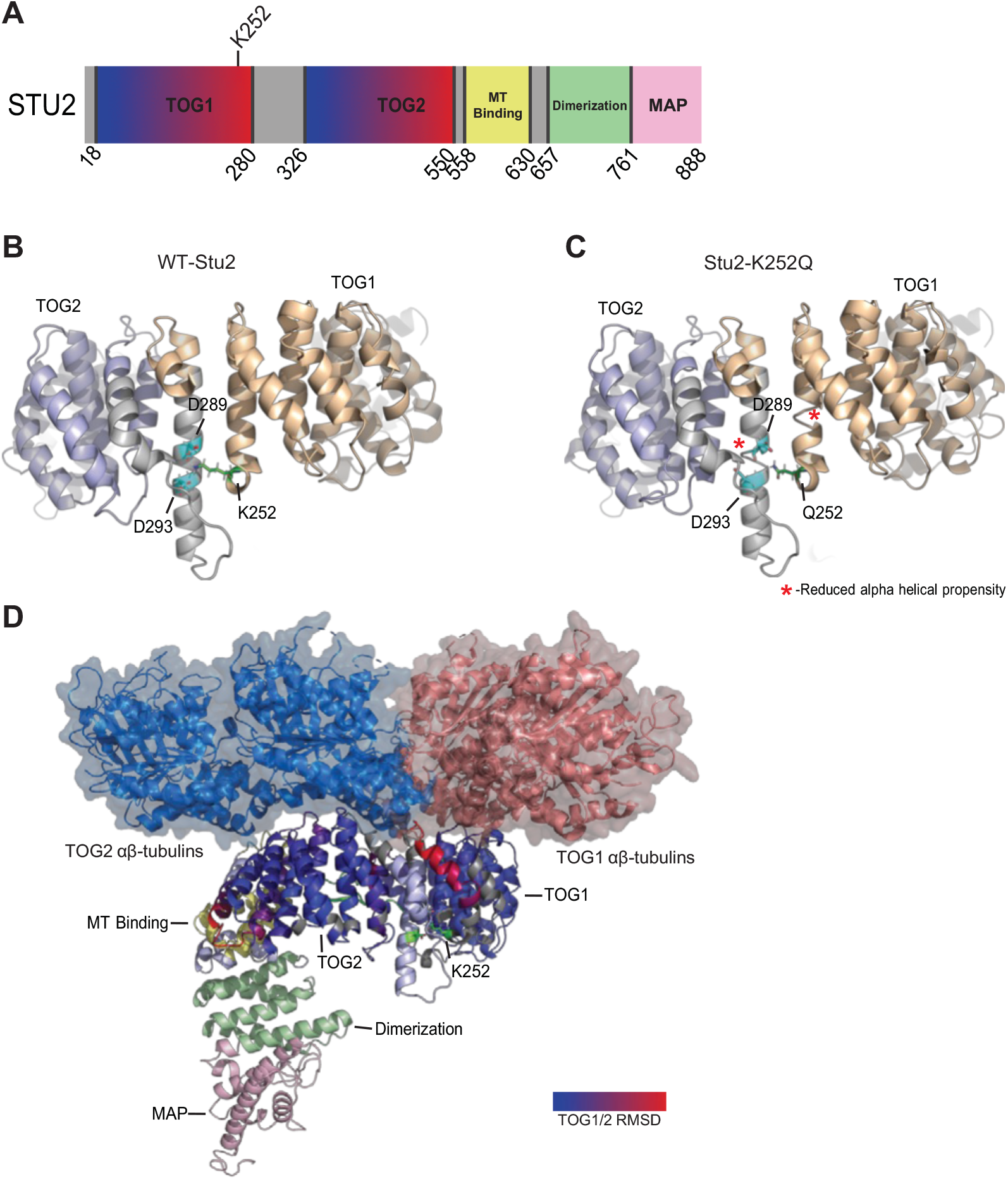
Modeling predicts that acetylation at Stu2 K252 might regulate intra-helical salt bridges between TOG1 and TOG2 domains. (A) Lysine 252 lies at the c-terminal end of TOG1. I-TASSER modeling of WT-Stu2 (B) and Stu2-K252Q (C) indicate that K252 forms salt bridges with D289 and D293 on an adjacent α-helix. This model predicts that these salt bridges confer stability to the intra-TOG domain linker region. We note that mapping the TOG1 section of the TOG1/ αβ-tubulin crystal structure (pdb:4FFB) and the TOG2 section of the TOG2/ αβ-tubulin crystal structure (pdb:4U3J) structure to our I-TASSER model resulted in a steric clash between the two coordinated tubulin heterodimers (D). This suggests that the I-TASSER model positions the two TOG domains too closely. Alternatively, we speculate that acetylation of K252 may increase the distance between the TOG domains. The two αβ-tubulin heterodimers are shown in blue and pink respectively.

In addition to the three lysines identified in this report, previous acetylomic studies have identified other acetylated lysines in XMAP215 proteins. For these however, confirmatory and functional analyses remain unexplored. Acetylation of *S. cerevisiae* K711 was detected in the acetylome work by Downey *et al*., 2015. This is consistent with our finding that additional acetylation sites should be present in Stu2 (Figure 3). Notably, K711 is not conserved in fungal systems (Figure S5A). Further, acetylation of the human XMAP215 family members CKAP5/ch-TOG were previously detected in two separate high-throughput proteomics screens (Svinkina et al., 2015) (Choudhary et al., 2009). Svinkina *et al*., identified five additional acetylation sites in human CKAP5 at positions K163, K1411, K1463, K1761, and K1769 (Svinkina et al., 2015). Choudhary *et al*. identified K48 in the human ChTOG / CKAP5 protein, (Choudhary et al., 2009), however, it is conserved only in higher eukaryotes, but not in fungi. In *Rattus norvegicus*, Lundby et al., 2012 (Lundby et al., 2012) identified a peptide consisting of amino acids D733-K743 in which either K741 or K743 was acetylated. Notably, in human the acetylation site at K163 and the rat acetylation site at K741 lie on the analogous alpha helices of their respective TOG domains. Together, this supports the notion that the functional importance of Stu2 acetylation may be extendable to other systems.

Acetylation has been shown to regulate several other microtubule associated proteins. For instance, acetylation of K212 in the human plus-end tracking protein EB1 (the homologue of the yeast Bim1) was shown to regulate interactions with CLIP-170, p150^glued^, and APC (Xie et al., 2018). Additionally, EB1 acetylation at K220 plays a role in EB1 localization to plus ends of mitotic microtubules and helps coordinate the timing of metaphase alignment (Xia et al., 2012). Acetylation of CLIP-170 (the homologue of yeast Bik1) was found to influence cellular migration of pancreatic cancer cells (Li et al., 2014). Lastly, acetylation of Ndc80 residues K53 and K59 by TIP60 is important for mitotic chromosome transmission (Zhao et al., 2019). Together with our Stu2 data, these findings suggest that acetylation may be a general mechanism by which MAPs are regulated.

In summary, this work proposes a novel hypothesis for the regulation of Stu2 association with kinetochores, spindle pole bodies, and microtubule plus ends. As the expression status of the human XMAP215 family member, CKAP5/ch-TOG is associated with a poorer prognosis in cancers (Charrasse et al., 1998), future work is necessary to understand how acetylation affects other XMAP215 family members, particularly in disease states.

## MATERIALS AND METHODS

### Detection of Stu2 acetylated lysines using mass spectrometry

Yeast expressing Stu2-HA (yRM10637) or a vector control (yRM10641) were grown to mid-exponential phase and disrupted using cryo-milling as described in Greenlee *et al*., 2018. Frozen powdered cell debris, termed “fluff,” was resuspended in 1x PBS containing 0.1% Tween, 20 mM N-Ethylmaleimide, 20 mM 2-Iodoacetamide, 1:500 Sigma protease inhibitor cocktail #8849, and 1 mM PMSF, and clarified as described in Greenlee *et. al*., 2018. Protein concentrations were determined by Bradford assay using BSA for the standard curve.

Resuspended extract (75 mgs) was incubated with anti-HA magnetic beads for 1 hr at 4 °C and washed 3 times with 1x PBS containing 0.1% Tween. To elute, the beads were boiled in 100 µL of 2.5X Laemmli sample buffer for 5 min. Proteins were resolved on 10% SDS PAGE until 50 kDa markers were run off the gel to ensure optimal band separation. The SDS-PAGE gel was then fixed in 50% methanol, 10% acetic acid for 1 hr and stained for 15 min using Coomassie blue. To confirm the presence of Stu2 in bands, 10% of each pull-down was run on an SDS- PAGE, transferred to a nitrocellulose membrane, and blocked overnight in 0.1% I-block reagent (Applied Biosystems, Bedford, MA) dissolved in PBS containing 0.1% Tween. The membrane was probed with mouse anti-HA to identify HA-tagged Stu2. SDS PAGE bands containing Stu2 were cut from the gel, extracted with acetonitrile, and digested with trypsin in 2M Urea. Notably, this analysis was performed using a wild-type yeast strain, with neither TSA nor genetic approaches, to detect Stu2 acetylation, reducing the potential that identified acetylation sites are experimentally induced.

Peptides from the digested samples were dissolved in 0.1% aqueous formic acid, and injected onto a 0.075 x 400 mm nano HPLC column packed with 3-µm Magic AQ C18 particles. Peptides were separated using a 120-min gradient of 3-30% acetonitrile/0.1%formic acid and eluted through a stainless-steel emitter for ionization in a Proxeon ion source. Peptide ions were analyzed by a high/high mass accuracy approach. Parent ions were measured using the Orbitrap sector of a Fusion mass spectrometer (Thermo), followed by data-dependent quadrupole selection of individual precursors and fragmentation in both the ion trap (CID at 35% energy) and in the HCD cell (25% or 35% energy). Lastly, fragmented ions were measured again in the Orbitrap sector at 30,000 resolution.

Raw mass spectra files were converted to .mgf files using msconvert from Proteowizard (Chambers et al., 2012). SearchGUI was then used to analyze .mgf files with the proteomics search engines Comet, MS Amanda, MS-GF+, MyriMatch, OMSSA, and X!Tandem simultaneously (Barsnes & Vaudel, 2018). Fixed modifications in the searches included methionine oxidation, cysteine carbamidomethylation, and lysine carbamylation and variable modifications included lysine and N-terminal acetylation and phosphorylation of serine, threonine, and tyrosine. All searches omitted matches outside of a ±10 ppm mass accuracy window. SearchGUI outputs were then combined using PeptideShaker to compare peptide identifications from each independent search engine.

### Detection of AcK-Stu2 bands by western blotting

For western blotting analysis, the lysine deacetylase inhibitor Trichostatin A (TSA) was used to preserve Stu2 acetylation for subsequent detection. Yeast cells were grown to an OD_600_ value of 0.3 and treated with a final concentration of 200 ng/mL TSA in DMSO or DMSO alone for 2 hr at 30 °C. Following TSA treatments, cells were harvested, cryo-milled, resuspended in 1x PBS containing 0.1% Tween, 20 mM N-Ethylmaleimide, 20 mM 2-Iodoacetamide, 1:500 Sigma protease inhibitor cocktail #8849, and 1 mM PMSF, and clarified as described in Greenlee *et. al*., 2018. Protein concentrations were determined by Bradford assay using a BSA standard curve. Whole cell extracts containing 91 mgs of protein were prepared from strains expressing Stu2-HA (yRM12358), untagged Stu2 (yRM12359), Stu2-3KR-HA (yRM12364), or Stu2-3KQ-HA (yRM12369), as described in Table 1. These were incubated with anti-HA magnetic beads (Pierce Biotechnology Inc, and Thermo Fisher, Waltham, MA), washed, and eluted with 100 uL of 2.5x Laemmli sample buffer, and boiled for 5 min. 20% of each sample was resolved by SDS-PAGE gels and transferred to nitrocellulose membranes for immunoblotting with anti-HA (Santa Cruz, clone F-7) to detect Stu2 or anti-AcK (Genetex, Acetyl Lysine antibody clone 1C6) to detect acetylation. In these analyses, we found that the speed with which the extracts were processed and immunoblotted was the most important factor in achieving blots with detectable acetylated Stu2.

### Conservation among Stu2, Tub2, Tub4 homologues

To determine the conservation of lysines shown to be acetylated, we compiled a list of twenty-eight XMAP215 family members from twenty-seven different organisms including: *S. cerevisiae* (P46675), *M. sympodialis* (A0A1M8A6U5), *D. discoideum* (Q1ZXQ8), *P. pallidum* (D3BU59), *A. deanei* (S9W6U7), *P. marinus* (C5LPC0), *C. elegans* (G5EEM5), *T. conorhini* (A0A3R7L7I8), *E. multilocularis* (A0A087W255), *A. thaliana* (Q94FN2), *S. scitamineum* (A0A0F7S5S5), *O. sativa subsp. Japonica* (Q5N749), *P. humanus subsp. Corporis* (E0VU40), *A. darlingi* (W5J5U5), *A. aegypti* (A0A1S4FBN3), *D. melanogaster* (Q9VEZ3), *X. laevis* (Q9PT63), *M. musculus* (A2AGT5), *H. sapiens* (Q14008), *S. pombe* (Q94534, Q09933), *A. gossypii* (Q75CQ1), *P. antarctica* (A0A081CMB9), *G. candidum* (A0A0J9XCS4), *C. albicans* (A0A1D8PTZ8), *C. glabrata* (A0A0W0D3T8), *E. nidulans* (Q5B1Q9), and *U. maydis* (A0A0D1CA73). These MT polymerases were grouped as either fungal organisms or broadly as eukaryotes, and aligned using Clustal Omega (Sievers & Higgins, 2018).

Conservation analysis of β- and γ-tubulins was carried out using Clustal Omega as previously described using β-tubulins from *S. cerevisiae* (P02557), *C. elegans* (P12456), *A. thaliana* (P12411), *D. melanogaster* (Q24560), *X. tropicalis* (Q6GLE7), *M. musculus* (A2AQ07), and *H. sapiens* (P07437) and γ-tubulins from *S. cerevisiae* (P53378), *C. elegans* (P34475), *A. thaliana* (P38557), *D. melanogaster* (P23257), *X. tropicalis* (P23330), *M. musculus* (P83887), and *H. sapiens* (P23258).

### Complementation of *STU2* deletion

To evaluate the viability of Stu2 acetyl mutants, we performed a plasmid shuffle assay. Cultures (5 mL) of yeast containing a *URA3*-marked plasmid expressing wild-type *STU2* (pRM10693) and a *LEU2-*marked plasmids expressing *STU2* (yRM12337), vector (yRM12338), K252R (yRM12340), K469R (yRM2341), K870R (yRM12342), 2KR (yRM12343), 3KR (yRM12344), K252Q (yRM12345), K469Q (yRM12346), K870Q (yRM12347), 2KQ (yRM12348), or 3KQ (yRM12349) were grown overnight. OD600 values were determined for each culture and normalized to 0.100 each. Normalized cultures were further diluted to prepare 0.100, 0.055, and 0.010 dilutions in 96 well plates. These were transferred with a multi-pronged device to synthetic defined media lacking histidine, leucine, and uracil to confirm equal loading of cells. Cells were transferred to 5-FOA plates lacking leucine to select for cells that lost the WT-*STU2 URA3+* plasmid, while retaining the Stu2 version on the LEU2 plasmid.

While dissecting *STU2*/*stu2*-Δ1::*HIS3* heterozygous diploid strain (yRM 11105) for subsequent yeast mating assays, we noticed that the Stu2-3KR mutant failed to produce viable spores in the wild-type *STU2* background. Of the 36 tetrads dissected from multiple independent transformations, 72 spores were expected to contain the Stu-3KR plasmids in the wild-type *STU2* background. However, zero viable spores of this genotype were obtained (Table S2). The fact that Stu2 can function as a dimer leads us to speculate that acetylation of Stu2 might play some role in regulating microtubule function during germination.

### Benomyl resistance of Stu2 mutants

Cultures of yeast containing *STU2* or the acetylation mutants were normalized to OD600 values of 0.100 and diluted as described above. Cells were transferred to leucine deficient media to confirm equal loading and plates lacking histidine, leucine, and uracil to confirm the absence of the *URA3* WT-*STU2* plasmid. To test the sensitivity of Stu2 mutant to microtubule stress, cells were also transferred to DMSO, DMSO + 10 μg/mL of benomyl, or 20 μg/mL of benomyl and grown at 23, 30, and 37 °C. To ascertain whether observed phenotypes are dominant, the Stu2 acetyl-lysine mutants were also tested for benomyl sensitivity in a *STU2*/*stu2Δ* heterologous diploid.

### Strain construction for chromosome loss assay: chromosome III fragmentation

To investigate Stu2’s role kinetochore function, we generated a diploid yeast strain containing an artificial 125-kb chromosome fragment expressing the sup11 suppressor, as described in (Shero et al., 1991). This assay requires *ade2* mutation in the background, which was generated by mating the *MAT*a strain yRM11408 and the *MAT*α strain yRM11407) for 1 hr. The resulting zygotes were isolated by micromanipulation using an Olympus BX41 dissection microscope. The resulting diploid (yRM11988) was heterozygous for *STU2*/*stu2Δ* and homozygous for the *ade2-101*. The artificial chromosome was generated by the fragmentation of Chromosome III, induced through the integration of NotI linearized pJS2/pRM11972 plasmid. (Figure S3A,B).

Following the creation of the artificial chromosome, colonies were selected based on a pale-pink color phenotype associated with presence of single chromosome fragments. To detect the presence of the artificial chromosome, we electrophoretically karyotyped the candidate strains using pulsed field gel electrophoresis (PFGE). Successful candidates were identified by comparing them to a previously published haploid cell-line possessing the 125-kb chromosome III fragment (YCTF58 / yRM11971 from (Kroll et al., 1996)) as a positive control, and to the parental “no chromosome fragment” strain (yRM11988) as a negative control. For this analysis, whole chromosome samples were prepared and visualized using methods adapted from (Carle & Olson, 1985) and (Warren et al., 2002). To create the necessary yeast protoplasts, yeast cultures (5mLs) were collected at 3000 RPM for 3 min using an Allegra X-15R Beckman Coulter tabletop centrifuge. Growth media was discarded, cell pellets were washed 3 times with 1.6 mLs of 50 mM EDTA, pH 7.5. Cells were resuspended in 300 µL of 50 mM EDTA, pH 7.5 to which 500 µL of 1% low-gelling-temperature agarose (Sigma Aldrich, St. Louis MO) (prepared in 125mM EDTA, ph7.5) was added. Then, 100ul of digestion buffer containing 1 mg/mL zymolyase, 5% β-mercaptoethanol, in SCE (1M sorbitol, 100mM sodium citrate, pH 8.0) at 37 degrees was added. Protoplast mixes were transferred to 24 well plates and allowed to solidify at room temperature. After solidification, 500 µL of a liquid overlay layer containing 270 mM EDTA –pH 9.0, 10 mM Tris –pH 8.0, and 7.5% β-mercaptoethanol was added to each cell suspension and incubated overnight at 37 °C in a sealed plastic bag.

To proteolytically digest the protoplasts, the liquid overlay was removed and replaced with a solution containing 1 mg/mL proteinase K resuspended in 270 mM EDTA (pH 9.0), 10 mg/mL N-Lauroylsarcosine (pH 9.0), and 10 mM Tris (pH 8.0). Protoplasts with proteolysis buffer were incubated overnight at 50 °C in a sealed plastic bag. To store digested cells, the proteolysis liquid overlay was removed and replaced with 500 µLs of pH 9.0 0.5 M EDTA.

Chromosome III fragmentation events, illustrated in Figure S3C, were screened using PFGE in a BIORAD CHEF-DR III system. Gel slices of each digestion were transferred to 1.7 mM wells of a 1% agarose gel made with pH 8.0 0.5x TBE buffer containing 45 mM Tris-borate and 1 mM EDTA. PFGE was carried out in 0.5x TBE buffer at 200 V and 14 °C for 30 hr using continuous ramp switch times of 24 to 54 seconds. To visualize bands, the gel was stained in 0.5 µg/mL ethidium bromide in ddH_2_O for 1 hr and destained in ddH_2_O for 1.5 hr. Using electrophoretic mobility to separate the four smallest chromosomes in *S. cerevisiae*, we identified two strains with Chromosome III fragmentation (Figure S6D.). One of the strains was heterozygous for Chr III fragmentation and possessed an unaltered Chr III along with greater (Chr III GF) and lesser fragments (Chr III LF) from sister Chr III fragmentation (yRM12011). The second strain was homozygous for full length Chr III but still possessed a band at the molecular weight range diagnostic of a Chr III fragment but lacked the Chr III greater fragment (yRM12012).

We note that any type of mechanical stress in the form of squeezing or bumping of the gel slices would increase the amount of chromosome shearing, lower the chromosome integrity and leading to smearing of bands on the gel. This seemed most apparent during the excision of the gel slice and its subsequent deposition into the running gel.

### Quantitative chromosome loss assay

To quantitatively determine the rates of chromosome transmission fidelity for the strains described above, cells were initially grown over night to OD600 values of 0.100 to 0.200. Cells were further diluted to OD600 values of 0.010 in chilled water or chilled water containing 1 mg/mL 5-FOA and stored at 4 °C for 4 hr. Chilled yeast were diluted 100 fold and approximately 100-200 cells (125 µL) were then transferred to four SD plates lacking both uracil and leucine. Approximately 17,000 Cells from the 5-FOA treated cells were plated to 3 1 mg/mL 5-FOA plates lacking leucine and grown for 4 days at 30 °C. To quantify chromosomal loss, initial cellular densities were back calculated using the uracil leucine deficient plates.

Colonies growing on 5-FOA plates were quantified as individual chromosomal loss events. The total number of 5-FOA colonies was then divided by the total number of colonies predicted to determine a quantitative value for chromosomal loss.

### Qualitative chromosomal loss assay

To determine relative rates of chromosome transmission fidelity between mutants and WT-Stu2, yeast strains containing *SUP11* artificial chromosomes and plasmids expressing WT- Stu2 (yRM12253), vector with endogenous Stu2 (yRM12300), K252R (yRM12261), K469R (yRM12301), K870R (yRM12271), 2KR (yRM12266), 3KR (yRM12269), K252Q (yRM12262), K469Q (yRM12260), K870Q (yRM12270), 2KQ (yRM12267), and 3KQ (yRM12268) were initially grown overnight to OD600 values between 0.100 and 0.200. Overnight cultures were further diluted to approximately 0.0001 OD600 values. 25 or 50 cells of each strain were plated onto SC deficient for leucine with either 4 µg/mL (20%) or 8 µg/mL (40%) of normal adenine concentrations. Each set of conditions was repeated three times to generate error bars. Yeast were then grown at 30 °C for 5 days and stored for 1 day at 4 °C. Because sectors slowly become more apparent at low temperatures, all visible red sectors were counted within eight hr of each other. For each plate, the total number of sectors relative to colonies was determined as qualitative values.

### Co-precipitation of αβ-tubulin heterodimer or γ-tubulin with Stu2

Yeast were grown to an OD600 value of 0.7, harvested, cryo-milled, and resuspended in either PEM buffer (80 mM PIPES, 1 mM EGTA, 1 mM MgCl2, pH 6.8) containing 20 mM N-Ethylmaleimide, 20 mM 2-Iodoacetamide, 1:500 dilution of protease inhibitor cocktail (Sigma Aldrich, catalog #8849), and 1 mM PMSF or 1x PBS (pH 7.4) containing 0.1% Tween, 20 mM N-Ethylmaleimide, 20 mM 2-Iodoacetamide, 1:500 dilution of protease inhibitor cocktail (Sigma Aldrich, catalog #8849), and 1 mM PMSF. Resuspended extracts were then clarified at 27,000 x g. To detect the interactions of α-tubulin, Stu2-HA was immunoprecipitated from 67 mg of yeast extracts resuspended in PEM buffer using 20ul with anti-HA magnetic beads (Pierce Inc, Waltham, MA, #88836 magnetic beads). To detect interactions with γ-tubulin, Stu2-HA was immunoprecipitated from 90 mg of yeast extracts resuspended in 1x PBS, as described above.

To detect co-precipitated tubulins, pull-downs were subsequently western blotted and probed with 1:100 mouse anti α-tubulin (4A1 developed by Dr. Margaret Fuller (Stanford University) as a gift from Dr. Jeff Moore (University of Colorado, Anschutz Medical Campus)) and 1:1000 rabbit anti γ-tubulin (clone 137822 Abcam. Inc, Boston, MA). Stu2 was detected using mouse anti-HA (F-7 clone, SC7392, Santa Cruz Biotechnology, Santa Cruz, CA). Secondary antibodies for these blots were goat anti-rabbit conjugated with HRP from Jackson Immuno Research Labs (cat #111-035-144) (1:5000) and goat anti-mouse-HRP (1:5000) (Jackson Immuno Research Labs, Inc. West Grove, PA cat #115-036-146).

### Structural modeling of β- and γ-tubulin interactions with the Stu2 TOG2 domain

Pymol was used to visualize atomic coordinate data from the following pdb files: TOG/GTP β-tubulin complex from 4U3J, GDP β-tubulin polymerized in MTs from 5W3F, and GDP hydrolyzed γ-tubulin from 3CB2. We used the ColorByRMSD Python Module (Shandilya, 2016) to align and color code positional conservation for a small highly structured region of β- and γ-tubulins that includes the K469 interacting residues (β-EFPD/γ-RYPK) and the GTP hydrolyzing E-site (exchange site) (Ayaz et al., 2014) (Howes et al., 2017) (Rice et al., 2008).

We next extracted α-tubulin data from the pdb:4U3J file and repeated this process to align β-tubulin residues with γ-tubulin residues from open (pdb:5FM1) and closed (pdb:5FLZ) conformations of γ-TuSC structures (Greenberg et al., 2016). This produced a structural model for how the Stu2 TOG2 domain would hypothetically interact with both γ-TuSC conformations.

### Stu2 structure prediction

The structural prediction of Stu2 was generated with I-TASSER (Yang et al., 2015) (Roy et al., 2010) (Zhang, 2008) using the full Stu2 sequence as a template (uniprot:P46675, SGD:S000004035). To determine the impact of K252Q mutations on Stu2 structure, the full amino acid sequence was submitted to I-TASSER with the K252Q amino acid substitution and cross-referenced with the WT K252 structure.

## ACKNOWLEDGEMENTS

This work was supported by grants from the NIH (R15GM119117-01) and the Oklahoma Agricultural Experiment Station (OKL02961) and USDA National Institute of Food and Agriculture (Hatch project #9377) to RKM. SM was supported in part by an CASNR/OAES Scholarship and a Niblack Research Scholarship, both at OSU.

## Abbreviations

AcK: acetyl-lysine
MAP: Microtubule associated protein
PBS: Phosphate buffered saline
PFGE: Pulsed field gel electrophoresis
SC: synthetic complete
SIM: SUMO interacting motif
SPB: spindle pole body
WT: wild type
YPD: yeast peptone dextrose
+TIPs: +Tip interacting proteins

## SUPPLEMENTAL FIGURES AND TABLES

**Figure S1.**
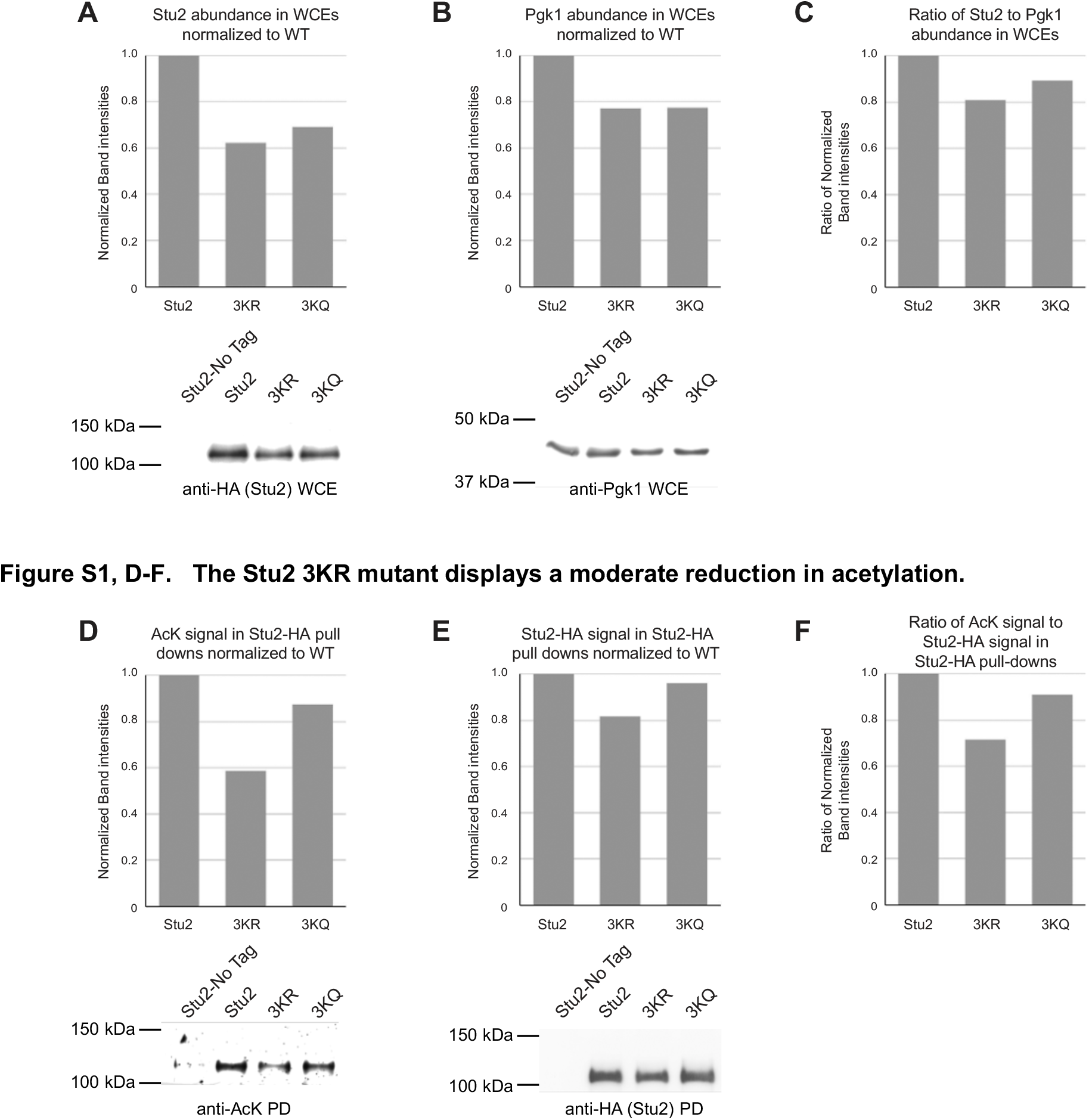
Stu2 triple mutants do not appreciably affect the steady state abundance of Stu2. Densitometry analysis of western blots probing (A) Stu2-HA and (B) Pgk1 from WCEs was performed to determine the (C) relative steady state abundances of Stu2 mutants. The Stu2 3KR mutant displays a moderate reduction in acetylation. Densitometry analysis of western blots probing (D) acetyl-lysine and (E) Stu2-HA from Stu2 immunoprecipitations was performed to determine the (F) reduction of Stu2 acetylation present in Stu2 3K mutants.

**Figure S2.**
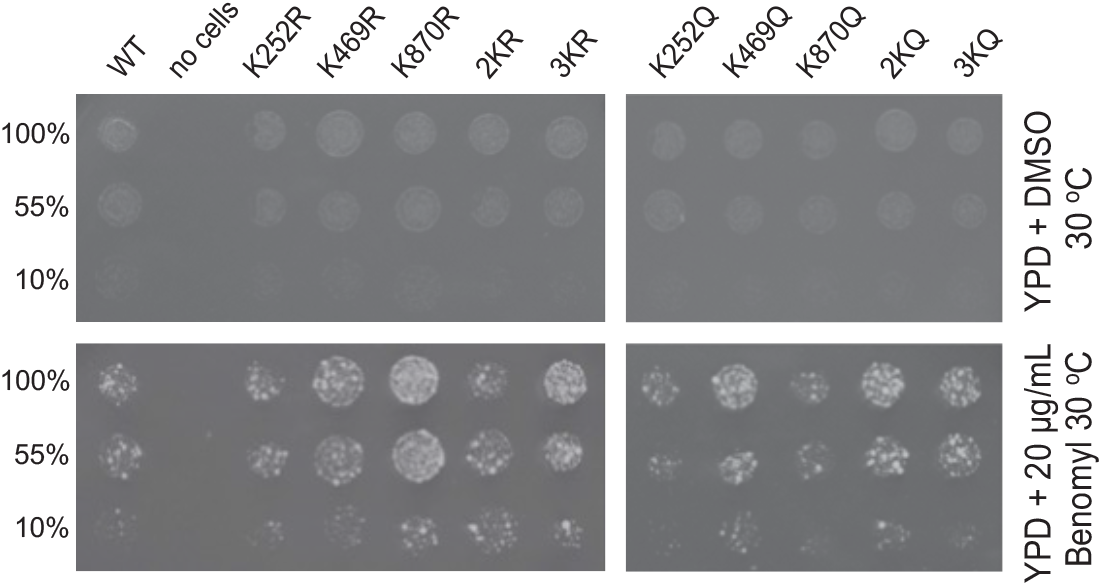
Stu2 acetylation states also confer resistance to microtubule-depolymerization stress at 30 °C. Yeast containing *LEU2* plasmids with *STU2* (yRM12358), K252R (yRM12360), K469R (12361), K870R (12362), 2KR (yRM12363), 3KR (yRM12364), K252Q (yRM12365), K469Q (yRM12366), K870Q (yRM12367), 2KQ (yRM12368), or 3KQ (yRM12369) were transferred to YPD containing DMSO or DMSO + 10 μg/mL benomyl and grown at 30 °C.

**Figure S3.**
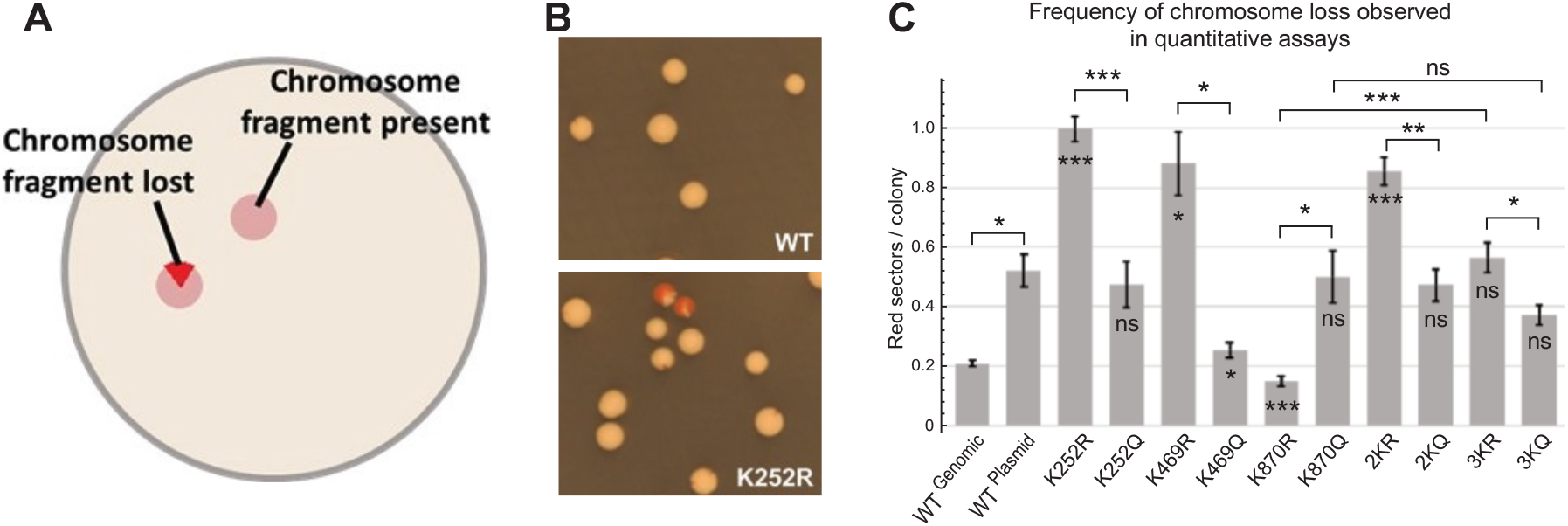
Qualitative chromosome loss assays indicate that Stu2 acetylation impacts chromosome distribution. (A) Chromosome transmission fidelity was qualitatively assessed using a color-sectoring colony assay in which the red phenotype is suppressed by *SUP11* present on the 125-kb artificial chromosome (pRM11972/pJS2). (B) Colonies with *STU2* plasmids (pRM2119) form red sectors less frequently than cells containing K252R plasmids (pRM11481). Yeast in this assay contained plasmids with *STU2* (yRM12253), K252R (yRM12261), K469R (yRM12301), K870R (yRM12271), 2KR (yRM12266), 3KR (yRM12269), K252Q (yRM12262), K469Q (yRM12260), K870Q (yRM12270), 2KQ (yRM12267), and 3KQ (yRM12268) in a *stu2Δ::HIS3* background, except for the WT-genomic control which had an intact *STU2* at its genetic locus and an empty vector instead of the *STU2* on a plasmid (yRM12300). (C) The total number of sectors counted was divided by the total number of colonies present to determine chromosomal loss rates of mutants relative to WT controls. Qualitative chromosomal loss data displayed as mean ± SEM of triplicate plates; * p < 0.05, ** p < 0.01, *** p < 0.005 (raw qualitative data are available in supplementary file 1).

**Figure S4.**
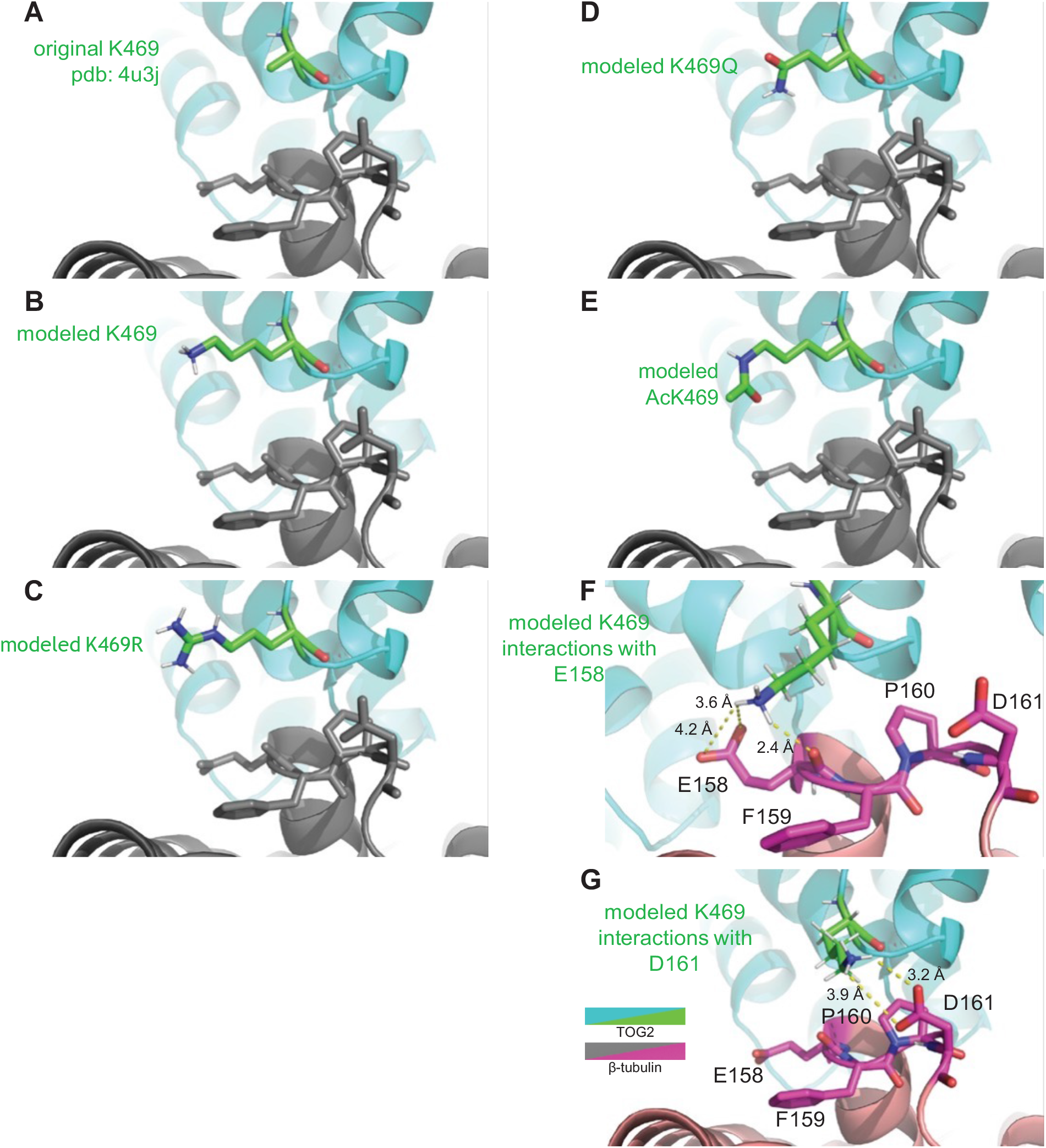
The side chain of K469 was not solved in crystal structures between TOG2 and tubulin heterodimer in pdb:4U3J (A). Here we modeled K469 using PyMOL to gain insight into how K469 may interact with β-tubulin. The TOG2 K469 side chain is reconstructed in panel B. We modeled side chains for the (C) K-to-R acetylation prevention mutation, the (D) K-to-Q acetylation mimetic mutation, and (E) acetylated K469. Lastly, K469 rotamer confirmations in this model were examined to further model this residue’s interactions with (F) E158 and (G) D161 of β-tubulin.

**Figure S5.**
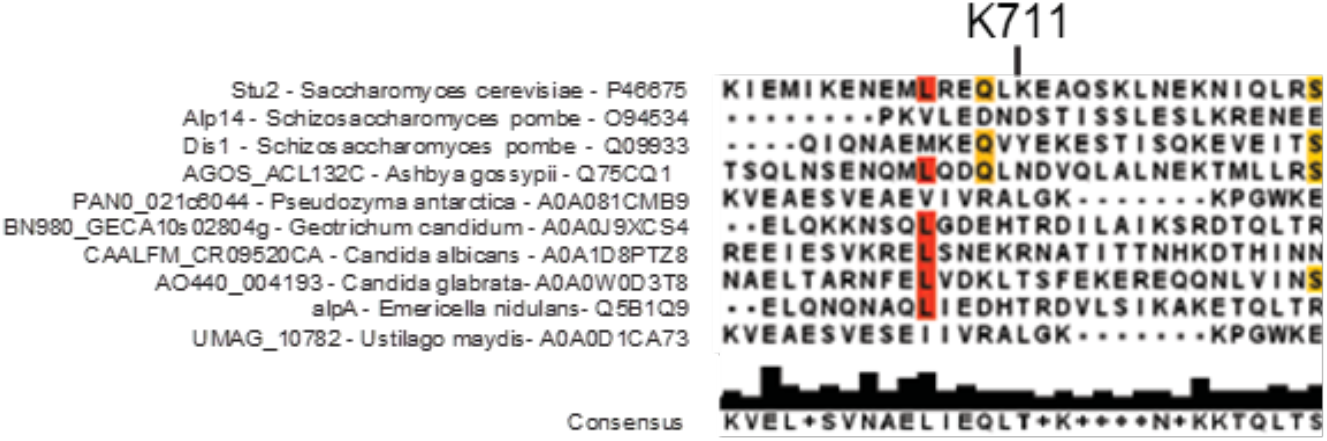
A previous acetylomic screen identified K711 as an acetylated lysine in the yeast XMAP215 family member, Stu2 (Downey et al., 2015). To understand its evolutionary conservation, we aligned the K711 with a series of homologues from several fungal species as described above in Figure 2. This analysis suggests that the K711 site is not evolutionarily conserved.

**Figure S6.**
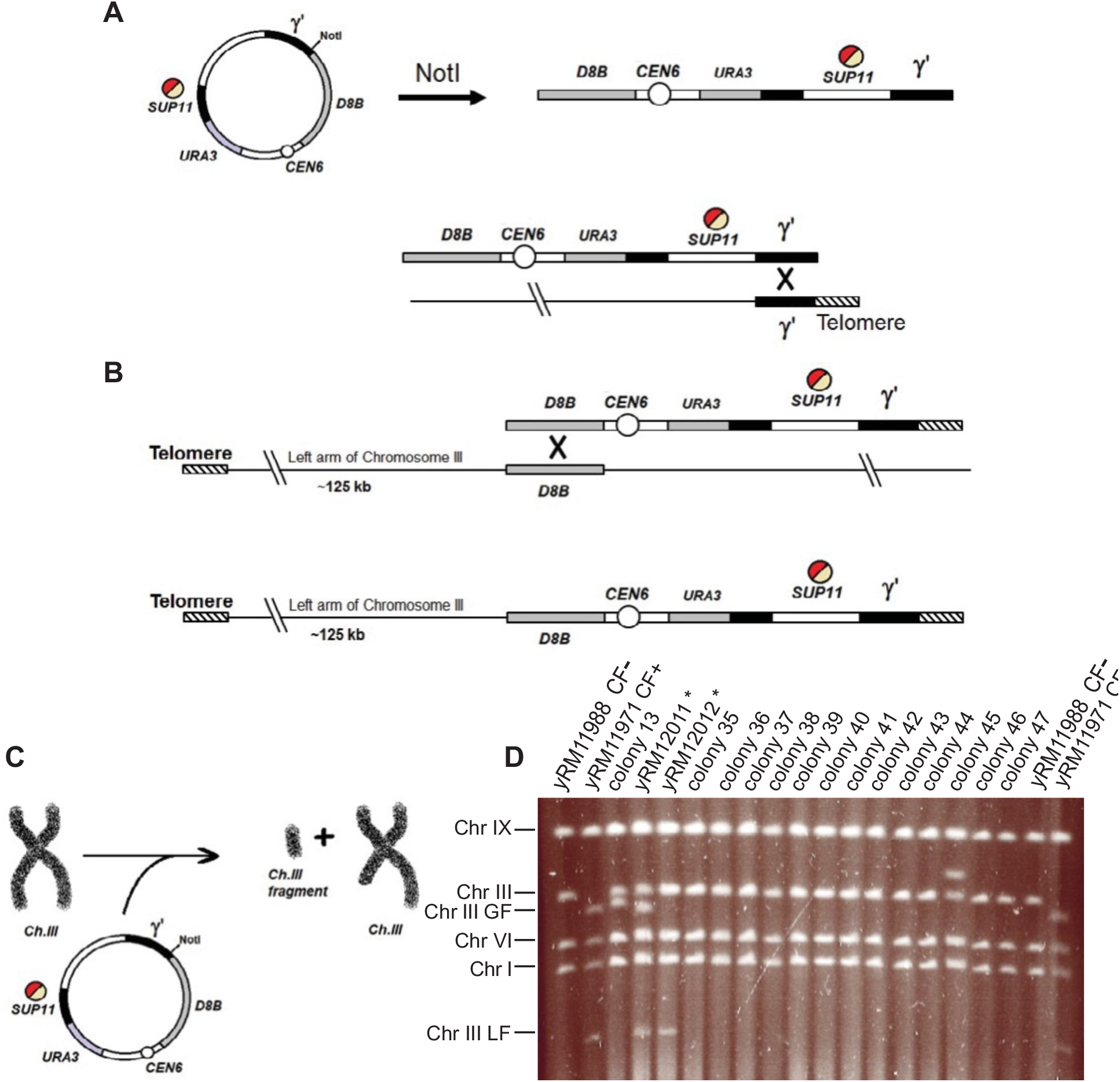
Fragmentation of Chromosome III adapted from (Shero et al., 1991). (A) Linearization and cross one during pJS2/pRM11972 integration occurs between the γ’ element of the vector and endogenous yeast γ’ sequences to generate a chromosome fragment precursor with a one sided telomeric DNA cap. (B) A second cross occurs in the D8B genetic sequence specific to the left arm of chromosome III to yield a 125-kb Chromosome III fragment. (C) A simplified model regarding the aim of chromosome fragmentation. (D) PFGE electrophoretic karyotyping of the four smallest chromosomes of S. cerevisiae, the generated Chromosome III greater fragment, and the 125-kb Chromosome III lesser fragment.

**Table S1.** All Stu2 AcK spectra detected. Additional mass spectra were identified for each acetylated residue. Listed are all of the confidence scores associated with each putative AcK specta. AcK 252 had a total of 32 spectra with the following confidence scores: 9 with 100%, 5 with 99%, 4 with 97%, 8 with 96%, 2 with 95%, 1 with 94%, 1 with 83%, 1 with 82%, 1 with 32%. AcK 469 had a total of 10 spectra with the following confidence scores: 1 with 97%, 3 with 96%, 1 with 95%, 1 with 94%, 1 with 93%, 1 with 91%, and 1 with 70%. AcK 870 had a total of 18 spectra with the following confidence scores: 3 with 100%, 4 with 99%, 3 with 98%, 3 with 97%, 1 with 96%, 1 with 86%, 1 with 85%, 1 with 82%, and 1 with 80%. All spectra fell well within a 30 ppm mass accuracy with the vast majority of parent ion masses deviating less than 3 ppm.

| AcK site | 100% confidence | 90-99% confidence | 80-89% confidence | <80% confidence | Total spectra |
| --- | --- | --- | --- | --- | --- |
| AcK 252 | 9 | 20 | 2 | 1 | 32 |
| AcK 469 | 0 | 8 | 0 | 1 | 9 |
| AcK 870 | 3 | 11 | 4 | 0 | 18 |

**Table S2.** Tetrad dissection summary. A *STU2*/*stu2-Δ1*::*HIS3* heterozygous diploid (yRM11105) was transformed with wild-type *STU2*-HA (pRM2119), empty vector (pRM2200), untagged wild-type *STU2* (pRM6507), K252R (pRM11481), K469R (pRM11249), K870R (pRM12016), 2KR (pRM11966), 3KR (pRM11976), K252Q (pRM11482), K469Q (pRM11254), K870Q (pRM12015), 2KQ (pRM11969), and 3KQ (pRM11974). Each strain was then sporulated and tetrads dissected. *STU2* deletion was marked with *HIS3* (*stu2*-Δ1::*HIS3*). Yeast centromeric plasmids expressing *STU2* or *stu2* mutants were marked with a *LEU2* gene. In this analysis, the total tetrads for each dissected strain are reported. The expected viable spores reflect the 2:2 nature of sister chromatid distribution as well as the inability of our vector control to rescue *stu2* deletion. Observed spores represents the total number of spores that grew. The four categories of possible genotypes are reported in columns 5- 8. Notably, the K252,469,870R (3KR) mutant fails to produce viable spores in the presence of an endogenous copy of *STU2*. In the absence of endogenous *STU2* (*stu2-Δ1::HIS3*), spores containing the 3KR mutant grow.

|  | Total<br>Tetrads | Expected<br>Viable<br>Spores | Observed<br>Spores | Percent<br>viability of<br>expected<br>spores | *stu2-*<br>*Δ1::HIS3+*<br>/ *leu2-* | *stu2-*<br>*Δ1::HIS3+*<br>/ *LEU2+* | *STU2*<br>(his-)<br>/ *leu-* | *STU2* (his-) /<br>*LEU2+* | Expected<br>*STU2+* (his-)<br>*LEU+* Spores | % of expected<br>*STU2+* (his-)<br>*LEU+* Spores<br>Observed |
| --- | --- | --- | --- | --- | --- | --- | --- | --- | --- | --- |
| Vector Ctrl | 156 | 312 | 285 | 91.3% | 0 | 0 | 190 | 95 | 312 | 30.4% |
| WT-HA | 158 | 632 | 417 | 66.0% | 126 | 126 | 174 | 117 | 316 | 37.0% |
| WT-NT | 28 | 112 | 60 | 53.6% | 12 | 12 | 33 | 15 | 56 | 26.8% |
| K252R | 36 | 144 | 119 | 82.6% | 50 | 50 | 16 | 53 | 72 | 73.6% |
| K469R | 48 | 192 | 138 | 71.9% | 48 | 48 | 46 | 44 | 96 | 45.8% |
| K870R | 12 | 48 | 39 | 81.3% | 16 | 16 | 9 | 14 | 24 | 58.3% |
| K252,469R | 47 | 188 | 128 | 68.1% | 40 | 40 | 38 | 50 | 94 | 53.2% |
| K252,469,870R | 36 | 144 | 55 | 38.2% | 27 | 27 | 28 | 0 | 72 | 0.0% |
| K252Q | 36 | 144 | 125 | 86.8% | 57 | 57 | 8 | 60 | 72 | 83.3% |
| K469Q | 24 | 96 | 62 | 64.6% | 23 | 23 | 16 | 23 | 48 | 47.9% |
| K870Q | 12 | 48 | 30 | 62.5% | 7 | 7 | 16 | 7 | 24 | 29.2% |
| K252,469Q | 48 | 192 | 130 | 67.7% | 40 | 40 | 50 | 40 | 96 | 41.7% |
| K252,469,870Q | 36 | 144 | 89 | 61.8% | 37 | 37 | 18 | 34 | 72 | 47.2% |

